# Cardioids reveal self-organizing principles of human cardiogenesis

**DOI:** 10.1101/2020.07.06.189431

**Authors:** Pablo Hofbauer, Stefan Jahnel, Nora Papai, Magdalena Giesshammer, Mirjam Penc, Katherina Tavernini, Nastasja Grdseloff, Christy Meledeth, Alison Deyett, Clara Schmidt, Claudia Ctortecka, Šejla Šalic, Maria Novatchkova, Sasha Mendjan

**Author notes:** These authors contributed equally to this work (Pablo Hofbauer, Stefan Jahnel, Nora Papai).

## Abstract

Organoids that self-organize into tissue-like structures have transformed our ability to model human development and disease. To date, all major organs can be mimicked using self-organizing organoids with the notable exception of the human heart. Here, we established self-organizing cardioids from human pluripotent stem cells that intrinsically specify, pattern and morph into chamber-like structures containing a cavity. Cardioid complexity can be controlled by signaling that instructs the separation of cardiomyocyte and endothelial layers, and by directing epicardial spreading, inward migration and differentiation. We find that cavity morphogenesis is governed by a mesodermal WNT-BMP signaling axis and requires its target HAND1, a transcription factor linked to human heart chamber cavity defects. In parallel, a WNT-VEGF axis coordinates myocardial self-organization with endothelial patterning and specification. Human cardioids represent a powerful platform to mechanistically dissect self-organization and congenital heart defects, serving as a foundation for future translational research.

**Highlights:**

- Cardioids form cardiac-like chambers with inner endothelial lining and interact with epicardium
- Cardioid self-organization and lineage complexity can be controlled by signaling
- WNT-BMP signaling directs cavity formation in self-organized cardioids via HAND1
- WNT-VEGF coordinate endothelial patterning with myocardial cavity morphogenesis

## INTRODUCTION

The human heart, the first functional organ to form in development, is one of the most difficult organs to model *in vitro* (Lancaster and Huch, 2019; Schutgens and Clevers, 2019). Malformations of the heart are by far the most common human birth defects (Majumdar et al., 2019) but their developmental etiology is poorly understood (Nees and Chung, 2019). These often dramatic morphogenetic disorders can be caused by mutations that affect the activity of cardiogenic signaling pathways and transcription factors during early embryogenesis (Kelly et al., 2014; Meilhac and Buckingham, 2010; Solloway and Harvey, 2003; Zaidi and Brueckner, 2017). From work in animal and cellular models we know how the cardiac lineage is specified in a stage-specific manner from embryonic mesoderm (Birket et al., 2015; Burridge et al., 2014; Costello et al., 2011; Lassar et al., 2003; 2001; Lee et al., 2017; Lian et al., 2012; Mendjan et al., 2014a) to produce cardiomyocytes (CMs), endocardial cells (ECs) and epicardial cells (Meilhac et al., 2014; Palpant et al., 2017). However, how signaling directs these cell types to self-organize into layers and shape a heart chamber, or fail in cardiac defects, remains unclear (Abu-Issa and Kirby, 2007; Kelly et al., 2014). These questions are challenging to tackle in complex systems, as manipulating signaling pathways at the spatial and temporal resolution required to dissect rapid and complex developmental processes is not yet feasible. Thus, we need *in vitro* models that mimic aspects of development but are simple enough to resolve the intricate dynamics of cardiogenesis and its malformations in humans.

Human organoids have been used to dissect mechanisms of patterning and morphogenesis in multiple tissues and organs. These structures emerge in culture from human pluripotent (hPSCs) or adult stem cells that are coaxed to form a tissue-like architecture, organ-specific cell types and exert organ-specific functions (Clevers, 2016; Lancaster and Knoblich, 2014) for use in numerous applications (Lancaster and Huch, 2019; Schutgens and Clevers, 2019). Their hallmark feature is the capability of self-organization that is driven by intrinsically coordinated specification, patterning and morphogenesis (Sasai et al., 2012; Sasai, 2013), in the absence of spatial constraints and interactions with other embryonic tissues. These properties allow the dissection of complex developmental processes in an organ-specific context as embryonic redundancies and compensation mechanisms are removed (Little and Combes, 2019; Sasai, 2013). For instance, pioneering work using PSC-derived optic cup organoids elucidated the morphogenesis of the eye (Eiraku et al., 2011; Nakano et al., 2012), gut organoids have been used to tease apart the initiation of intestinal crypt morphogenesis (Serra et al., 2019), and hPSC-derived brain organoids allow to study the etiology of human brain malformations (Lancaster et al., 2013; Qian et al., 2016). By harnessing developmental mechanisms, self-organization allows not only to study organogenesis and its defects, but results in more physiological models for a much wider range of diseases and applications (Lancaster and Huch, 2019; Tuveson and Clevers, 2019). Although self-organizing organoids have been reported for almost all major organs, there are currently no self-organizing human cardiac organoids that autonomously pattern and morph into an *in vivo*-like structure (Lancaster and Huch, 2019; Schutgens and Clevers, 2019).

Bioengineering approaches have successfully been applied to create artificially engineered heart tissues (often termed heart/cardiac organoids, microchambers, etc.) using scaffolds, molds, geometric confinement and protein matrices (Ma et al., 2018; Mills et al., 2019; 2017; Ronaldson-Bouchard et al., 2018; Tiburcy et al., 2017; Zhao et al., 2019). These have proven immensely useful to measure contraction force, perform compound screens and model structural muscle and arrhythmogenic disorders. Similarly, mouse and human PSC-derived 3D cardiac models including spherical aggregates (microtissues) of CMs and other cardiac cell types (Giacomelli et al., 2017; 2020; Richards et al., 2020), have been reported as promising tools for drug discovery; and embryoid models (Rossi et al., 2019; Silva et al., 2020) as providing insights into germ layer interactions in early organogenesis. However, existing models do not recapitulate cardiac-specific self-organizing patterning and morphogenesis to acquire *in vivo*-like architecture, and they are therefore limited as models of early human cardiogenesis and congenital heart disease.

Here, we established hPSC-derived self-organizing “cardioids” that undergo patterning and morphogenesis to form a cavity in the absence of non-cardiac tissues and exogenous extracellular matrix (ECM). Cardioids pattern into separate myocardial and endothelial layers and interact with migrating and differentiating epicardium, mimicking early heart chamber development. Using this system, we found that cardiac mesoderm self-organization and cavity formation is controlled by coordinated WNT- and BMP-signaling; such that absence of BMP target and transcription factor HAND1 results in a cardiac cavity defect that can be rescued by WNT signaling. We also found that cavity morphogenesis and endothelial patterning are coordinated by a mesodermal WNT and VEGF signaling at the cardiac mesoderm stage. Thus, human cardioids recapitulate myocardial, endothelial and epicardial morphogenesis and provide a powerful system to study self-organizing mechanisms of human cardiogenesis and congenital heart disease.

## RESULTS

### Formation of a cardiac chamber-like structure *in vitro*

To investigate whether an *in vitro* 3D chamber-like structure can be created intrinsically, we developed a differentiation approach based on temporal control of the key cardiogenic signaling pathways − Activin, BMP, FGF, retinoic acid and WNT. By recapitulating *in vivo* developmental stagin*g*, we sequentially specified hPSCs into mesoderm, cardiac mesoderm and beating cardiomyocyte progenitors at above 90% efficiency in 2D culture (Mendjan et al., 2014b) (Figure 1A). To screen for factors that are sufficient to stimulate intrinsic 3D cardiac structure formation in 2D culture (Figure S1A), we supplemented the media with selected ECM proteins that are involved in mesoderm development (Yap et al., 2019). Addition of Laminins 521/511 before mesoderm induction resulted in intrinsic self-assembly of cells and the striking formation of hollow, beating 3D structures expressing the CM marker TNNT2 after 7 days of differentiation (e.g. Figure S1A; Supplemental Video 1). When we performed cardiac differentiation entirely in 3D non-adherent culture, we found that exogenous ECM was not required for rapid self-assembly into cavity-containing CM structures (Figure S1B). Adaptation to a high-throughput differentiation approach allowed us to rapidly generate highly reproducible self-assembling cardiac 3D structures with cavities, hereafter referred to as cardioids (Figure 1B,C,D); Supplemental Videos 2 and 8).

**Figure 1.**
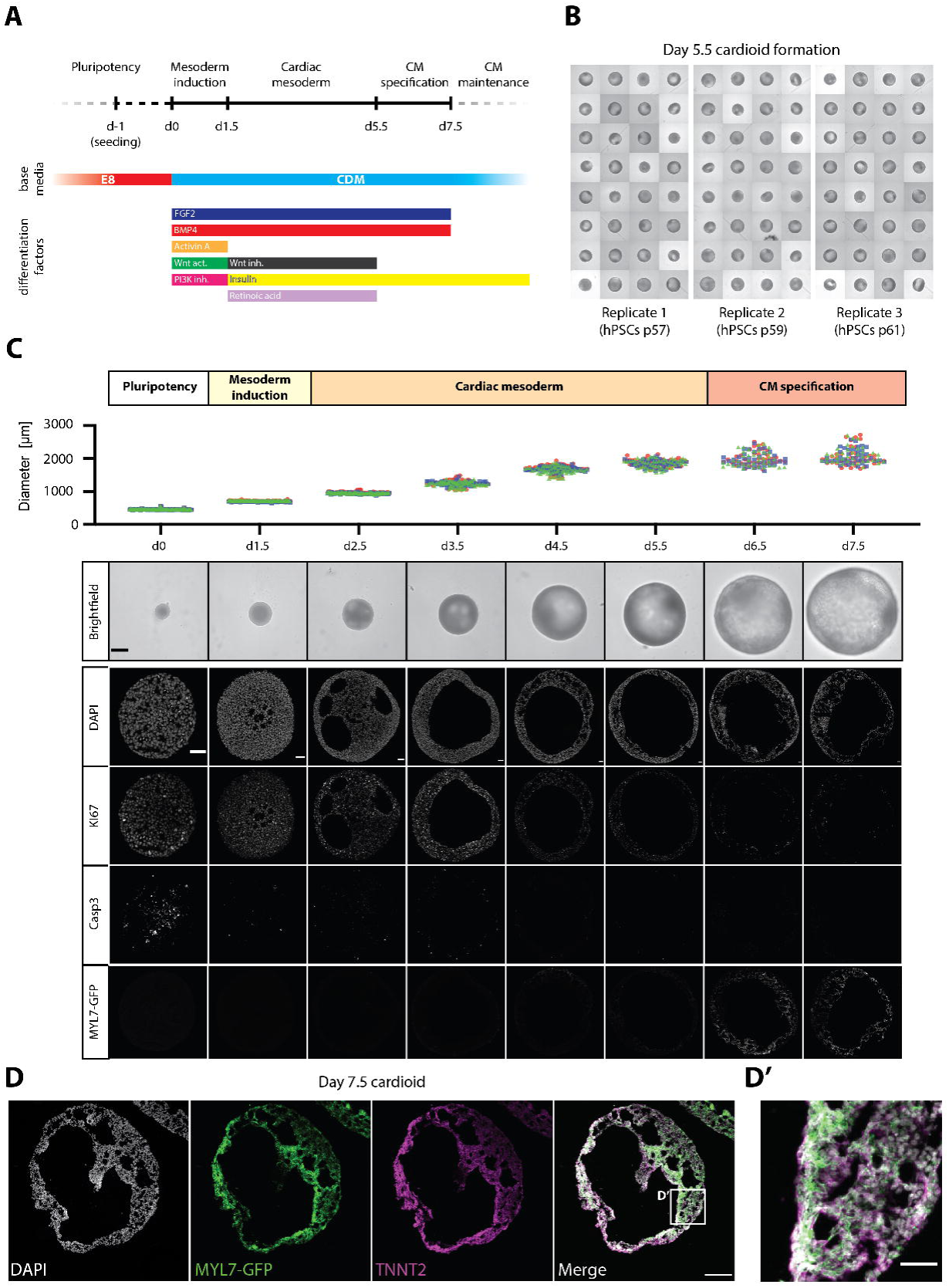
Formation of Cardiac Chamber-like Structures *in vitro*. **(A)** Cardiac differentiation protocol. WNT signaling activation (by CHIR99021), WNT signaling inhibition (by IWP-2/IWR-1/XAV-939), PI3K signaling inhibition (by LY294002). **(B)** Representative whole mount image (day 5.5) of a high-throughput differentiation approach showing robust generation of cavity-containing, beating structures. **(C)** CM differentiation time-course. Top: Quantification of size-change over time. Bottom: Whole mount brightfield images and sections showing the size increase and cavity formation over time. Cavities can first be seen at day 2.5 and are not formed by apoptosis (CASPASE3 negative) or regional proliferation (Ki67+) differences. Scalebars: 500μm (brightfield images), 50μm (sections). For quantification, the different symbols represent individual samples derived from three biological replicates. **(D)** Section showing the expression of the CM-specific markers MYL7 and TNNT2. Scalebar: 200μm. **(D’)** Detail of D. Scalebar: 50μm.

We next sought to compare properties of the cardioids relative to CMs differentiated in 2D. In both models, beating started between day 5 and 7 of differentiation at a similar rate and frequency (Ca^2+^ transients, beating frequency) (Figure S1C,D; Supplemental Video 3), and could be maintained in culture for at least three months (Supplemental Video 4). RNA-seq time-course analysis revealed an expression signature most similar to the first heart field lineage of cardiac mesoderm (HAND1^+^, TBX5^+^, NKX2-5^+^, TBX1^−^) (Figure 2A), which *in vivo* gives rise to the heart tube − the precursor of the left ventricular chamber and primitive atrium (Kelly et al., 2014). Genes encoding ion channels, structural proteins, cardiac transcription factors and sarcoplasmic reticulum proteins showed significantly higher expression levels in 3D cavity-forming structures, suggesting improved functionality (Figure S2A). GO-term analysis showed that cardioids exhibited gene expression patterns of heart morphogenesis and development, which were significantly upregulated over 2D CMs (Figure S2A,B) and aggregated 3D CM microtissues (Figure S2A,C). Overall, we successfully generated functional hPSC-derived cardioids that reproducibly self-assemble and can be maintained in long-term culture.

**Figure 2.**
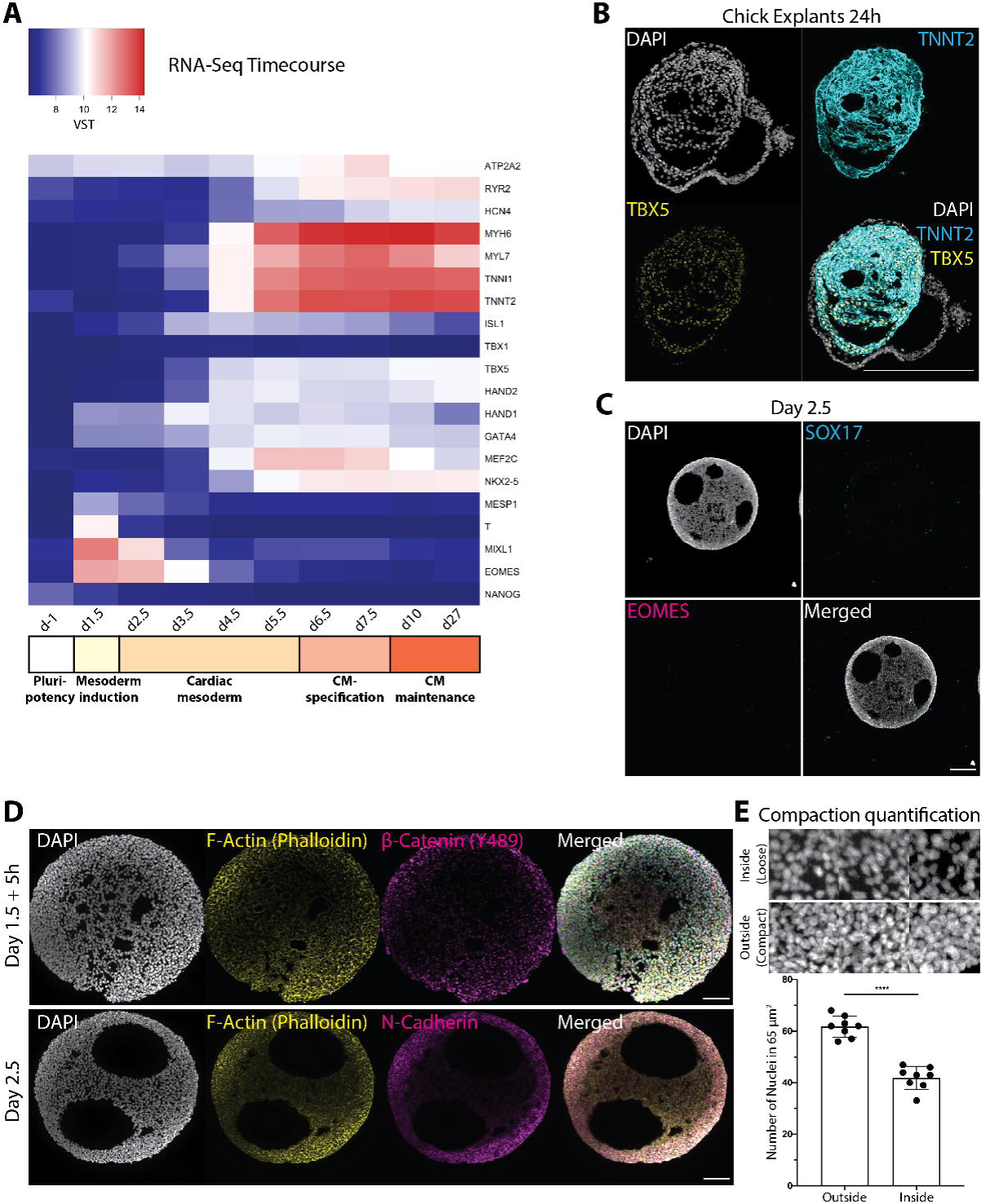
Cardiac Mesoderm Self-organizes Forming a Cavity *in vitro* and *ex vivo*. **(A)** Heatmap showing transcriptional change of key cardiac genes, including cardiac mesoderm markers between day 1.5 and day 5.5, during CM differentiation. VST: variance-stabilized transformed counts. **(B)** Cardiac mesoderm explants from chick embryos in human Cardiac Mesoderm conditions form chamber-like CM structures with cavities. Scalebar: 200 µm. **(C)** Self-organizing cardiac mesoderm at day 2.5 showing absence of SOX17+ and EOMES+ endoderm. **(D)** Cardiac mesoderm at day 1.5 (+5h) and at day 2.5 showing nuclear beta-CATENIN (phosphorylated Y489)) localization in the outer (compact) layer. **(E)** Images and quantification of number of nuclei in a 65 µm^2^ square in loose (inner) vs. compact (outer) layer of cardiac mesoderm. N=2.

### Cardiac mesoderm self-organizes forming a cavity *in vitro* and *ex vivo*

We next employed this system to ask whether the cavity within cardioids is formed by an intrinsic process of self-morphogenesis (Sasai, 2013) during specification in the absence of non-mesodermal tissues. By analyzing the time-course of cavity formation using live imaging and cryosectioning, we found that cavities were initiated and expanded robustly during the cardiac mesoderm (HAND1+) stage preceding expression of key cardiac structural markers, such as MYL7 (Figure 1C; Figure 2A, Supplemental Video 7). Most smaller cavities eventually coalesced into one major cavity (Figure 1C). Cavity expansion was not driven by either apoptosis or regional proliferation differences, as evidenced by cleaved CASPASE3 and KI67 staining (Figure 1C). Importantly, SOX17^+^/EOMES^+^ endoderm was absent during differentiation, indicating that cardiac mesoderm cavities were not generated as a result of endoderm instruction (Figure 2C). This is consistent with findings *in vivo*, as bilateral hearts can form when foregut endoderm morphogenesis is disrupted (Kuo et al., 1997; Li et al., 2004). Lumen formation also occurred when VEGF-driven endothelial cell (EC) differentiation was inhibited using the potent VEGFR inhibitor Sunitinib (Figure S2D). We concluded that cardiac mesoderm in our cultures is sufficient to intrinsically self-assemble into a CM-made cavity.

*In vivo*, cardiac mesoderm does not require foregut endoderm for basic morphogenesis of the heart in the mouse (Li et al., 2004) and in the chick (DeHaan and DeHaan, 1959). We therefore asked whether *ex vivo* dissected mesoderm from developing chick embryos can also form chamber-like structures in conditions we developed for human cardiac self-organization. Strikingly, in the absence of SOX2+ foregut, chick mesodermal explants developed into beating chamber-like structures *in vitro* similar to human cardiac mesoderm, demonstrating robust conservation of *in vitro* cardiogenic self-morphogenesis under permissive conditions (Figure 2B, Figure S2E,F).

Besides self-morphogenesis during specification, intrinsic self-patterning of a homogeneous starting cell population is a key hallmark of self-organization (Sasai, 2013). We therefore performed a time-course cryosectioning, staining and live imaging analysis of mesoderm and cardiac mesoderm to determine when the first self-patterning event occurs. While mesoderm at the induction stage appeared homogeneous, we observed differential localization of MYH10-GFP at the onset of the cardiac mesoderm stage (Figure 2D, Supplemental Video 5). Subsequent cavitation coincided with compaction at the periphery, reflected by higher cell density and accumulation of F-ACTIN, MYH10-GFP and N-CADHERIN (Figure 2D,E, Supplemental Video 5). It is likely that this compact cardiac mesoderm layer acts as a permeability barrier, as the cavity structures were impermeable to low-molecular-weight (4kDa) dextrans (Supplemental Video 6). In contrast, the inner part of the developing structures, where cavities first appeared, had less compact appearance with decreased N-CADHERIN signal (Figure 2D,E). These observations are consistent with the *in vivo* pattern of N-Cadherin in the compacted dorsal region of cardiac (splanchnic) mesoderm and the less compact region facing endocardial tubes and foregut endoderm (Linask, 2003). This far, we concluded that human cardioids feature the key hallmarks of self-organization (Sasai, 2013) – ongoing specification, intrinsic self-patterning into mesoderm layers and self-morphogenesis to shape a cavity.

### WNT and BMP control cardioid self-organization

We next used cardioids to dissect how signaling controls intrinsic morphogenesis and patterning during cardioid specification. To quantify phenotypes with high statistical power, we combined the high-throughput cardioid platform with a custom-made semi-automated imaging/analysis FIJI-pipeline. Using this setup, we examined which signals control cardioid self-organization and at what stage of mesodermal specification they act. We first systematically tested the effects of key mesoderm and cardiac mesoderm signaling dosages (e. g. WNT, BMP) on cardiac cavity self-morphogenesis. Surprisingly, we found that higher dosages of WNT signaling during mesoderm induction drove cavity expansion during the later cardiac mesoderm stage (Figure 3A,B), which has not been reported before. An intermediate WNT dosage promoted both cavity morphogenesis and CM specification. The optimal WNT activation range was consistent for each hPSC line but differed across lines, consistent with numerous studies showing line-specific signaling responses (Ortmann and Vallier, 2017; Strano et al., 2020). Importantly, the highest WNT dosage promoted cavity formation without CM specification (Figure 3A), highlighting a striking difference in signaling control of cell-fate specification versus morphogenesis.

**Figure 3.**
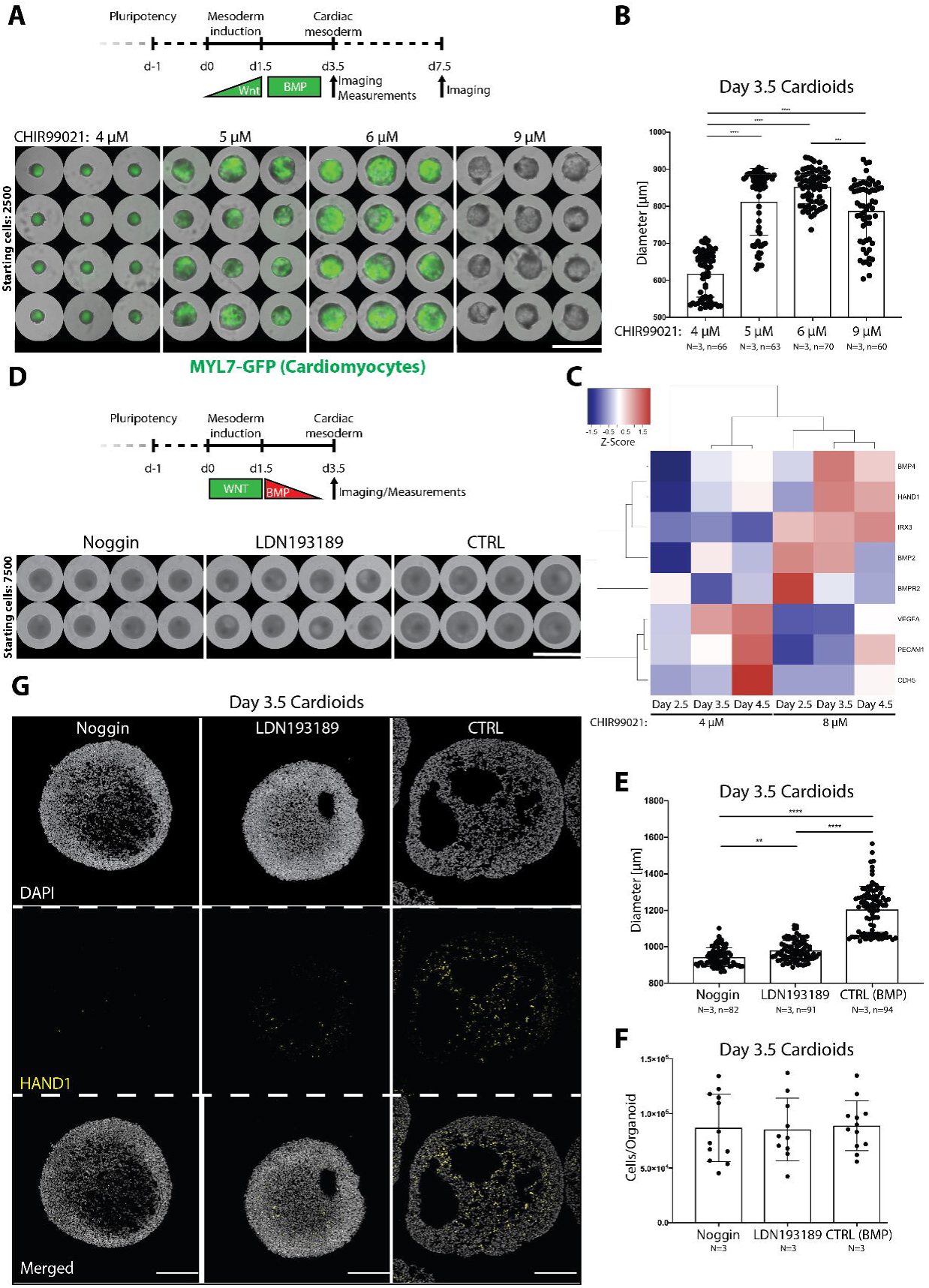
WNT and BMP Control Cardioid Self-organization. **(A)** Range of CHIR99021 concentrations during mesoderm induction shows dramatic effects on cardioid diameter (day 3.5) and CM specification (day 7.5). Protocol starting with 2500 hPSCs. Scalebar: 2500 μm. **(B)** Quantification of cardioid diameters at day 3.5 during cardiac mesoderm specification. **(C)** BMP target genes (HAND1, IRX3, BMP4, BMP2, BMPR2) are upregulated in cavity-forming conditions (8 μM CHIR99201), while EC genes (PECAM1, CDH5, VEGFA) are upregulated when 4 μM CHIR99021 is used. **(D)** BMP-inhibition with either Noggin (100 ng/ml) or LDN193189 (0.2 µM) reduces cardioid diameter. Protocol starting with 7500 hPSCs. Scalebar: 1958 μm. **(E)** Quantification of cardioid diameters at day 3.5 with or without BMP-inhibition. **(F)** Cell counting of cells/organoid reveals that the reduced diameter is not a function of fewer cells. **(G)** Noggin and LDN193189 treatments interfere with cavity expansion. Scalebars: 200 μm. All bar graphs show: Mean +/- SD.

To identify downstream mediators of WNT that control cardiac cavity morphogenesis, we performed RNA-seq analysis and compared gene expression profiles of mesoderm induced by higher (large cavity) and lower (small cavity) WNT signaling dosages. Among differentially expressed genes at the later cardiac mesoderm stage, we identified known cardiac mediators of BMP signaling (BMP4, BMP2, BMPR2) and some of its mesodermal targets (HAND1, IRX3) (Figure 3C) (Abu-Issa and Kirby, 2007; Lints et al., 1993; Riley et al., 1998). BMP drives cardiac specification at multiple stages, and we asked whether BMPs can instruct patterning and morphogenesis to form a cardiac cavity. To answer this question, we blocked BMP signaling using either the natural inhibitor Noggin or the compound LDN193189 during the initial two days of the cardiac mesoderm stage. BMP inhibition resulted in impaired cavity morphogenesis resulting in decreased cardioid size while cell number per cardioid remained stable (Figure 3D,E,F). Interestingly, upon BMP inhibition, the compact cardiac mesoderm layer expanded while the less compact cavity-forming layer shrank (Figure 3G). In contrast, WNT inhibition at the cardiac mesoderm stage was not necessary for cavity formation, although it is known to be essential for cardiac specification (Lassar et al., 2003; Yang et al., 2008)(Figure S3A). These data emphasize that control of specification versus morphogenesis can be fundamentally different processes, and that a mesodermal WNT-BMP signaling axis controls self-patterning and self-morphogenesis – both key self-organizing processes.

### HAND1 is required for cardiac mesoderm self-organization

Mutations in signaling and downstream transcription factors affect heart tube and chamber development and cause severe human cardiac malformations (Nees and Chung, 2019). For instance, in Hypoplastic Left Heart Syndrome, the most severe congenital defect in humans, disrupted levels of the BMP-regulated genes NKX2-5 and HAND1 are associated with a severely reduced cardiac cavity within the left ventricular chamber (Grossfeld et al., 2019; Kobayashi et al., 2014; Vincentz et al., 2017). The earliest phenotype in mutant Nkx2-5 and Hand1 mice manifests as defects in early left ventricular chamber morphogenesis (Firulli et al., 1998; McFadden, 2004; Riley et al., 1998; Risebro et al., 2006; Robb et al., 1995), but the disease etiology and the underlying morphogenetic mechanism in humans are less clear. Here, we asked whether NKX2-5 and HAND1 were required to achieve intrinsic self-organization in the absence of non-cardiac tissues. We generated knock-out (KO) hPSC lines for either HAND1 or NKX2-5 (Figure S4A,C,D). In NKX2-5 KO lines, we did not detect any cavity formation defects at the cardiac mesoderm stage. This is in line with the delayed onset of NKX2-5 compared to HAND1 expression in human cardiac mesoderm (Figure 2A), but in contrast to the mouse where Nkx2-5 is genetically upstream of Hand1 (Biben and Harvey, 1997; Tanaka et al., 1999).

In contrast, we observed a clear defect in cardiac cavity self-organization and size at the cardiac mesoderm stage in HAND1 KO lines (Figure 4A-D, Figure S4E). This phenotype manifested as cardioids of smaller size and with smaller and fewer cavities. Consistent with the cavity reduction, the compacted cardiac mesoderm layer expanded in KO compared to WT cardioids. These defects were not caused by a difference in cell number per cardioid, as both KO and WT showed similar cell counts (Figure 4E). Importantly, despite these defects in patterning and morphogenesis of cardiac mesoderm, early CM specification was not affected in HAND1 KO cardioids (Figure S4B). This again underlines the crucial distinction between control of cell specification vs. tissue patterning and organ morphogenesis that cardioids allow to dissect in the context of a cardiac malformation.

**Figure 4.**
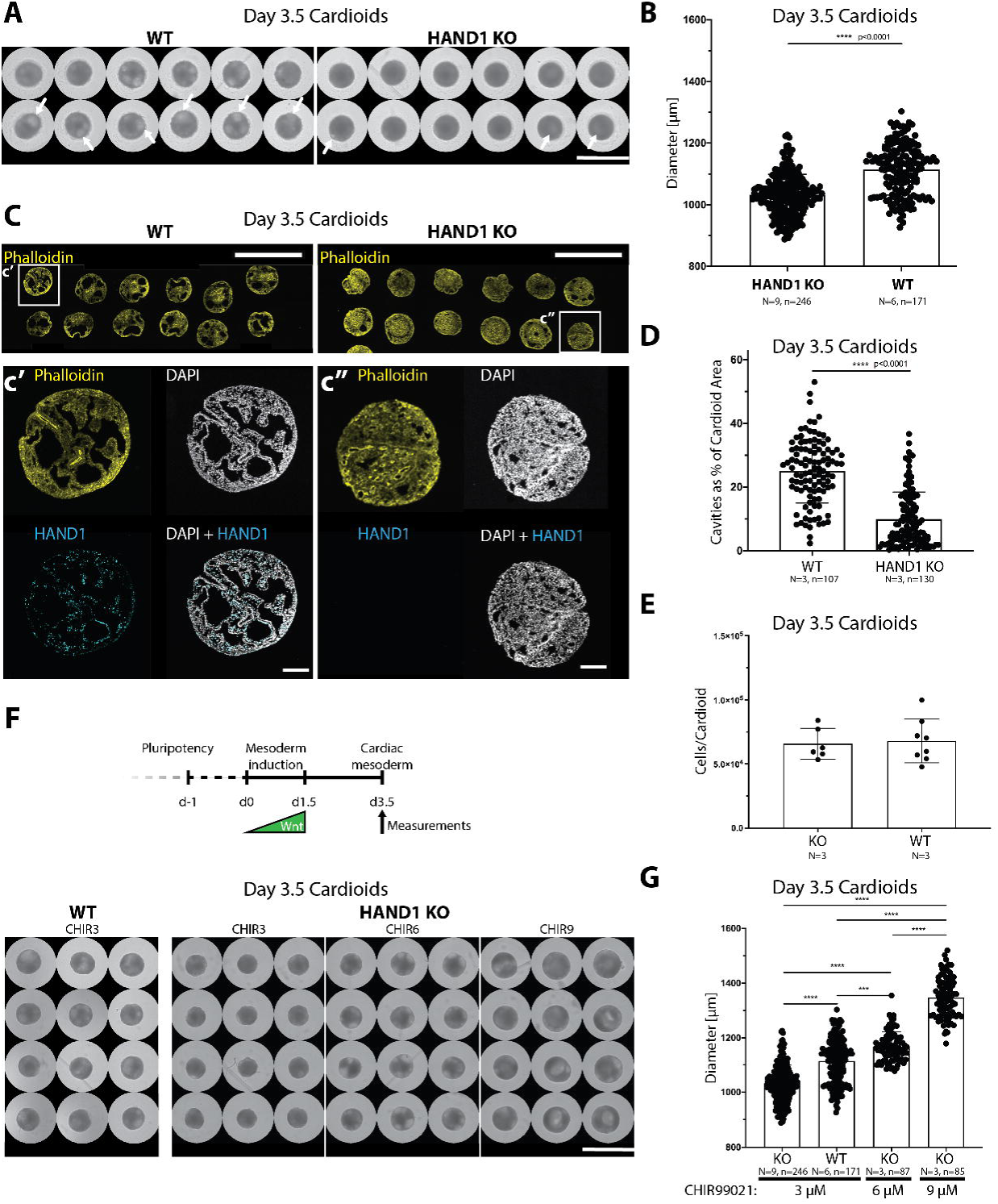
HAND1 is Required for Cardiac Mesoderm Self-organization. **(A)** WT cardioids show increased number of cavities (arrows) and diameter compared to HAND1 KO cardioids. Scalebar: 2000 μm. **(B)** Quantification of cardioid cavity expansion as a function of diameter in HAND1 KO and WT cardioids. **(C)** Phalloidin staining shows decreased number of cavities and their reduced size in HAND1 KO cardioids. Scalebars: 2000 μm. **(c’)** and **(c”)**, Detail of **C** and HAND1 KO confirmation by HAND1 staining. Scalebars: 200 μm. **(D)** Quantification of cardioid area covered by cavities in WT and KO cardioids. **(E)** Cell counting of cells/cardioid reveals that changes in diameter are not a function of cell number. **(F)** Timeline of organoid formation until day 3.5 (cardiac mesoderm stage), the point of analysis. Increased WNT signaling (CHIR99021) during mesoderm induction rescues lack of cavities in HAND1 KO organoids. Scalebar: 2500 μm. **(G)** Quantification of diameter shows that increased WNT activation rescues the HAND1 KO cardioid size defect. All bar graphs show: Mean +/- SD.

We next asked whether the HAND1 KO phenotype could be rescued by signaling. Increased dosage of WNT signaling during mesoderm induction rescued the HAND1 KO phenotype, confirming the role of WNT in cavity morphogenesis (Figure 4F,G). Consistently, HAND1 and IRX3, a human-specific early ventricular chamber marker (Cui et al., 2019), were induced in high WNT conditions that also promoted cavity expansion (Figure 3C). Moreover, HAND1 protein levels were diminished upon BMP inhibition, confirming that a WNT-BMP-HAND1 axis drives cardioid cavity self-organization (Figure 3G). Taken together, these data show that quantitative stage-specific signaling mechanisms of self-organization and genetic cardiac defects can be modeled in our high-throughput cardioid platform.

### WNT and VEGF coordinate endothelial and myocardial self-organization

We next explored whether cardioids can be used to dissect what signaling pathways program patterning and separation of the myocardium and endocardium to form an inner lining, a hallmark feature of the heart chamber. We discovered that a lower dosage of WNT activation during mesoderm induction stimulated spontaneous ECs differentiation within cardioids (Figure S5A), and that those ECs formed an inner lining of the cardioid cavity, resembling *in vivo* tissue structure. At the same time, as shown in Figure 3, a lower WNT activation dosage led to a smaller cardioid cavity. To probe these relationships, we compared an RNA-seq time-course of cardioids generated using high vs. low WNT activation dosage (Figure 5A) and found that VEGF-A was upregulated early in cardiac mesoderm by a lower WNT activation dosage. Other EC specifying genes (ETV2, TAL1, LMO2, PECAM1) were also upregulated (during the cardiac mesoderm stage) by the lower WNT activation dosage. In conclusion, an optimal WNT dosage during mesoderm induction simultaneously controls cavity self-morphogenesis, EC specification and the emergence of an endothelial lining of the cavity. We next asked whether VEGF signaling is sufficient to separate and pattern the CM and EC lineages in cardioids. To this end, we included VEGF-A after the CM specification stage and observed formation of an inner EC layer lining the cardioid cavity as *in vivo* (Figure 5B). To further elucidate whether we can control the self-organization of CM and EC layers in cardiac mesoderm, we included VEGF-A at this stage. We screened for optimal signaling conditions using a CM (MYL7-GFP) and EC (CDH5-Tomato) double-reporter hPSC line (Figure 5C,D,E, Figure S5B). As CMs and ECs co-differentiated from cardiac mesoderm into cavity-containing structures in the presence of VEGF-A, they separated into CM and EC layers with collagen I (EC) and fibronectin (CM and EC) expression, reminiscent of the *in vivo* situation (Figure 5D,E,G, Figure S5B). An intermediate WNT activation during mesoderm induction always resulted in an EC layer surrounding the CM layer (Figure 5E,G Figure S5B,D). This suggested that VEGF stimulates the early separation of the two layers, but that it is not sufficient to control the correct outer vs. inner orientation of the lining. In contrast, in vivo-like patterning of the EC layer and formation of an inner lining in the presence of VEGF at the cardiac mesoderm stage, could be achieved by lower WNT activation levels during mesoderm induction (Figure 5D). Thus, the dosage of WNT and timing of VEGF signaling coordinate specification, cavity morphogenesis and *in vivo*-like patterning of CM and EC lineages.

**Figure 5.**
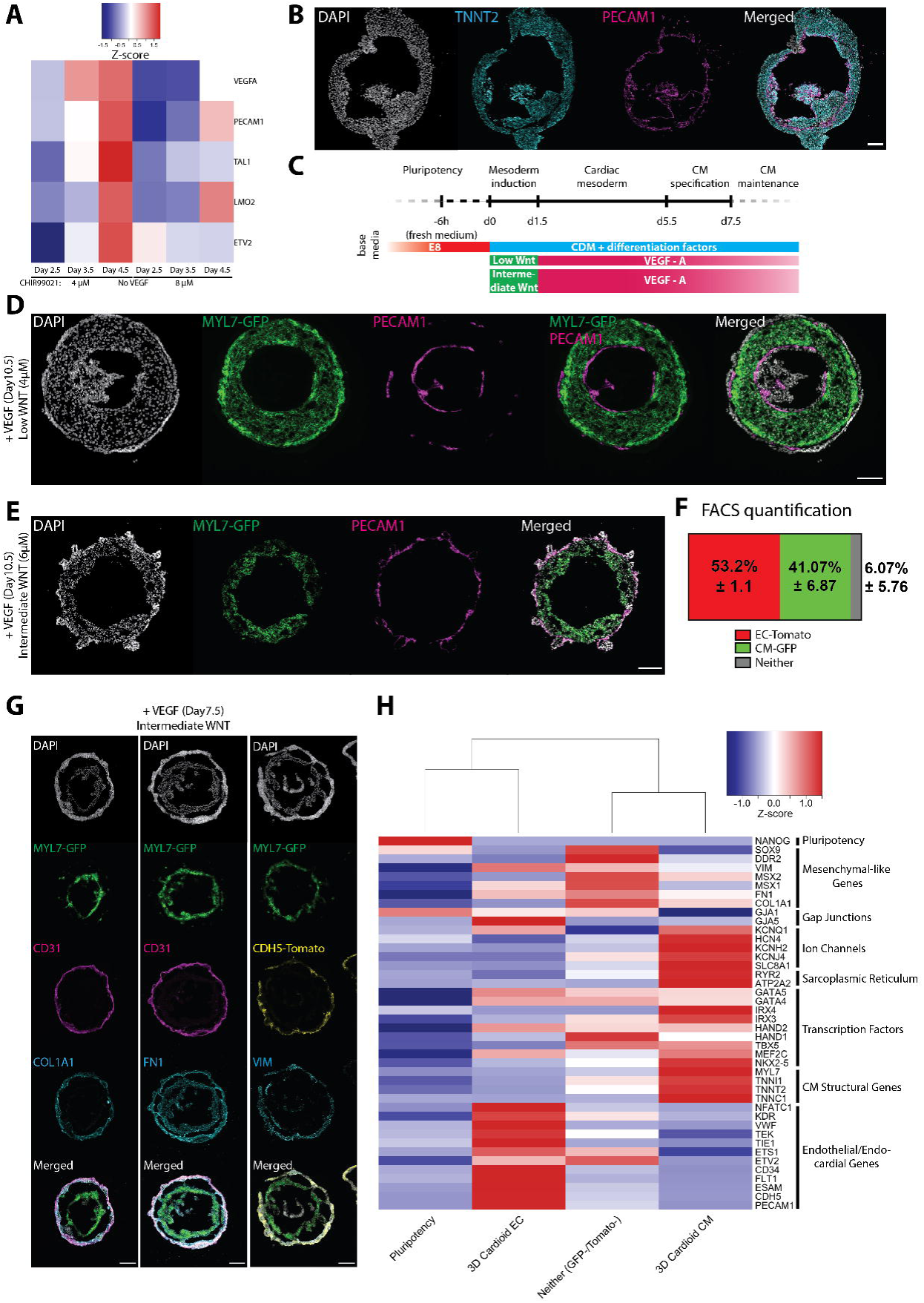
WNT and VEGF Coordinate Endothelial and Myocardial Self-organization. **(A)** Low WNT dosage (CHIR99021, 4µM) during mesoderm induction results in higher expression of EC-specifying genes (*VEGFA, TAL1, LMO2, ETV2, PECAM1*) compared to high WNT (CHIR99021, 8µM). **(B)** Addition of VEGF-A after CM specification for 7 days results in EC lining of the cavity and CMs. Scalebar: 100 µm. **(C)** Protocol to generate cardioids with entirely separate EC and CM layers by addition of VEGF-A from the cardiac mesoderm stage onward. **(D)** Low WNT dosage (CHIR99021, 4 µM) during mesoderm induction and VEGF-A treatment from cardiac mesoderm on results in inside EC lining a cavity and CMs. Scalebar: 100 µm. **(E)** Using protocol from **C** and with intermediate WNT dosage (CHIR99021, 6 µM) during mesoderm induction results in complete separation of EC and CM layers. Scalebar: 200 µm. **(F)** Quantification of FACS data showing a robust ratio of CMs and ECs in cardioids. Mean +/- SD. **(G)** Cardioids show separation of CM and EC layers as well as expression of ECM molecules COL1A1 (EC and Non-EC/Non-CM) and FN1 (all cells). Expression of fibroblast marker Vimentin by EC and Non-EC/Non-CM cells. Intermediate WNT dosage (CHIR99021, 6 µM) used. Scalebars: 200 µm. **(H)** Heatmap of Smart-Seq2 data of sorted hPSC, CM, EC, and Non-EC/Non-CM cells showing expression of the respective key genes. Non-EC/Non-CM (Neither) cells express genes related to putative EC-derived fibroblast-like cells.

To address whether CM and EC co-specification during cardiac mesoderm is necessary for self-organization, we developed control cardiac microtissues in which CMs and ECs are differentiated from cardiac mesoderm separately and then aggregated to form 3D structures (Figure S5C). In contrast to the co-differentiated cardioids, ECs formed intermingled networks within CMs, but they failed to pattern into layers. These data show that cardioids self-organize as they give rise to the first two major lineages of the heart.

### Endocardial-like identity of cardioid ECs

Although all self-organizing organoids share related tissue-like specification, patterning and morphogenesis processes, they are distinguished by their organ-specific cell types. We therefore next investigated the cellular heterogeneity within cardioids. Using intermediate WNT activation and VEGF, the ratio of CMs to ECs was remarkably stable at 41% (MYL7^+^) to 53% (CDH5^+^) (Figure 5F, Figure S5E,F), which was further corroborated by the deconvolution of bulk RNA-seq data using the MuSiC algorithm (Wang et al., 2019) and a cardiac cell-type-specific single-cell expression reference (Cui et al., 2019) for the developing human heart (Figure S6A). Smart-seq2 of sorted cells confirmed key CM and EC marker expression and GO-terms (Figure 5H, Figure 6C, Figure S6C), while the remaining 6% of the cells (MYL7^−^/CDH5^−^) expressed genes (e.g. SOX9, MSX1/2, COL1A1) related to putative EC-derived fibroblast-like cells (Huang et al., 2019; Neri et al., 2019) (Figure 5G,H, Figure S6D). No molecular signatures of either endodermal or ectodermal tissues were detected. Bulk proteomic analysis of cardioids confirmed their CM and EC proteome expression signature (Figure S6B).

**Figure 6.**
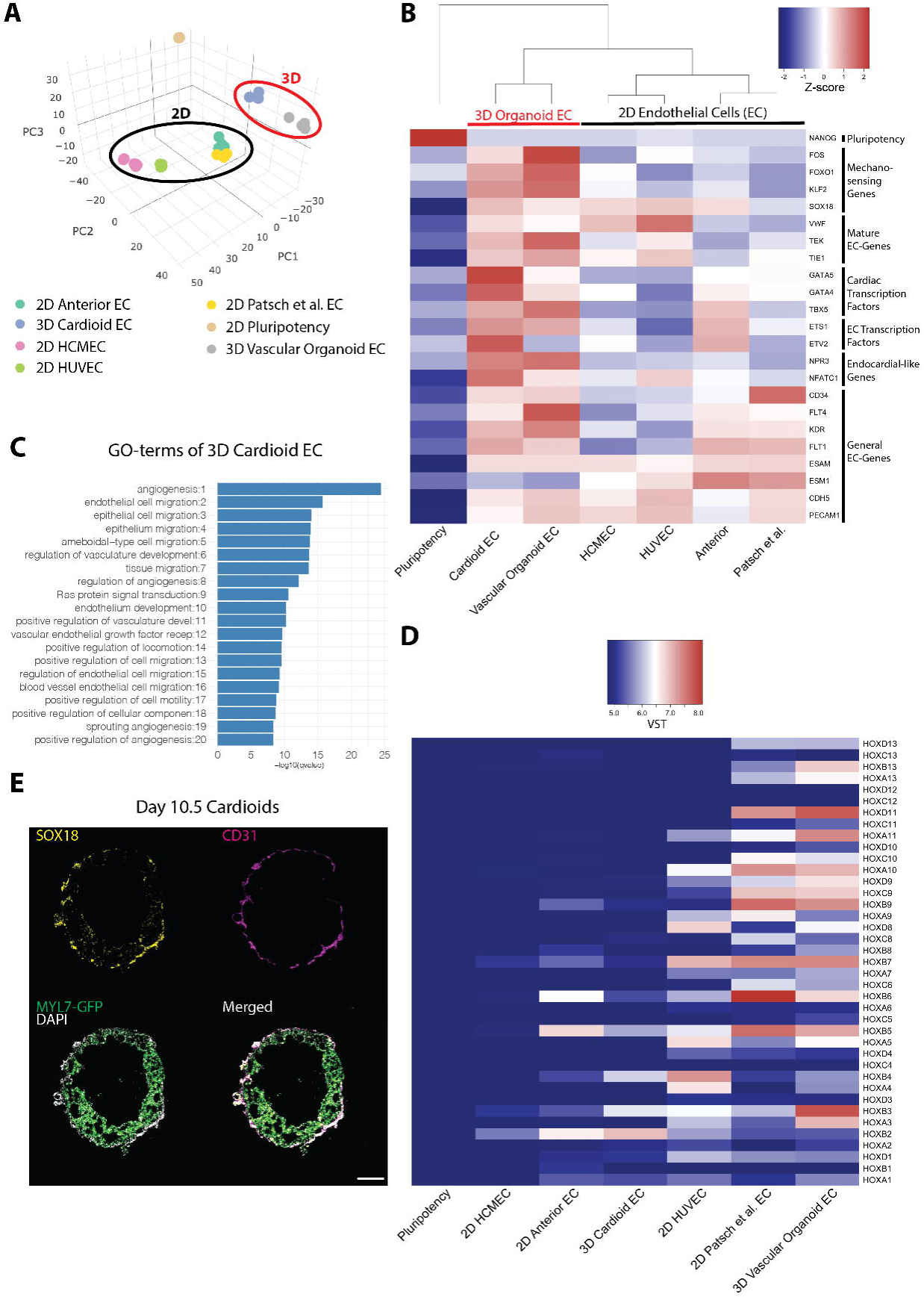
Endocardial-like Identity of Cardioid ECs. **(A)** PCA of: 2D-derived ECs (Anterior, Patsch et al., Human Cardiac Microvascular (HCMEC), Human Umbilical Vein (HUVEC)), hPSC and 3D-derived ECs from Cardioids (day 7.5, intermediate WNT dosage) and Vascular Organoids (Wimmer et al., Day>18)). **(B)** Differential expression of mechanosensing and maturation genes, cardiac transcription factors, EC transcription factors, endocardial-like and general EC genes in FACS-sorted ECs. **(C)** GO-terms of genes upregulated in cardiac ECs. **(D)** HOX-gene expression analysis of cardioid ECs, HCMECs and Anterior ECs shows expression of anterior HOX genes, while all other ECs show expression of posterior HOX genes as well. VST: variance-stabilized transformed counts. **(E)** SOX18 is expressed primarily in cardioid ECs. Scalebar: 200 µm.

All vascularized tissues and organs contain specific EC subtype, and accordingly, the endocardium has an identity-specific EC gene expression signature (Nakano et al., 2016). To determine EC identity in cardioids, we performed a Smart-seq2 analysis on sorted CDH5^+^ cardioid ECs and compared them to ECs generated using a well-established 2D differentiation protocol (Patsch et al., 2015), our 2D differentiation protocol using similar media conditions as in 3D, ECs from vascular organoids (Wimmer et al., 2019), human umbilical vein endothelial cells (HUVECs), and human cardiac microvascular endothelial cells (HCMECs) (Figure 6A,B). We found that cardioid-derived ECs were most similar to ECs from vascular organoids despite their age difference (Day 7.5 vs. Day >18). Importantly, ECs from cardioids showed increased transcript levels for cardiac transcription factors, such as GATA4/5, and genes associated with an endocardial-like identity (NFATC1, NPR3, (Tang et al., 2018)) (Figure 6B). Importantly, NFATC1 was also found in the proteomics data (Figure S6B). Their anterior HOX gene expression profile matched that of HCMECs from the adult human heart and that of ECs derived from cardiac mesoderm in 2D (Anterior ECs), but not that of the other analyzed more posterior EC subtypes (Figure 6D). Thus, the signature of ECs derived from cardioids is consistent with an endocardial-like identity.

The ability of endothelium to sense fluid flow, pressure and mechanical stretch is an instrumental requirement for its developmental and physiological roles, especially in the heart (Duchemin et al., 2019; Haack and Abdelilah-Seyfried, 2016). As expected for a bona fide model of cardiac development, a Smart-seq2 analysis of sorted ECs from cardioids revealed an upregulation of mechanosensitive genes (SOX18, KLF2, FOXO1, FOS) compared to ECs in 2D (Figure 6B,E), similar to the more matured 3D vascular organoid ECs. Markers of EC maturation (VWF, TEK, TIE1) were also upregulated in cardioids compared to 2D EC differentiations (Figure 6B). Overall, these results indicate that self-organization of CMs and ECs in 3D triggers essential aspects of endocardial identity and endothelial physiology.

### Epicardial interaction with cardioids

After inner endocardial-lined chamber formation follows outer epicardial engulfment of early myocardial chambers as the third major cardiac morphogenetic event (Cao et al., 2019; Simões and Riley, 2018). Guided by signals originating from the liver bud, a small clump of cells called the pro-epicardial organ develop into the epicardium which form the outer surface of the heart. (Andres-Delgado et al., 2019). After engulfment, signals from the CM layer (TGF-b, PDGF-b, FGFs) drive epicardial cell differentiation into smooth muscle cells (SMCs) and cardiac fibroblasts (CFs) – both crucial for further development and maturation of the heart (Cao and Poss, 2018). To study the intrinsic self-organization of this process in cardioids, we developed an epicardial differentiation protocol based on the signaling sequence known to specify the pro-epicardial organ in vertebrates (Figure 7A, Figure S8G) (Guadix et al., 2017; Iyer et al., 2015a; Witty et al., 2014). Importantly, the differentiation matched the estimated timeline of human epicardial development and was compatible with the cardioids in terms of timing and basic media conditions. Time-course of RNA-seq, immunostaining and flow cytometry analyses confirmed efficient epicardial differentiation in 2D and 3D as shown by expression of key epicardial markers (WT1, TCF21, TBX18) and early markers of differentiation (DCN, DDR2, COL1A1) (Figure S7A-E). As *in vivo*, depending on activation of TGF-b, FGF and PDGF signaling, pro-epicardial cells differentiated and efficiently induced the SMC marker ACTA2 and the CF marker COL1A1 (Figure S8A,B) and VIM (data not shown).

**Figure 7.**
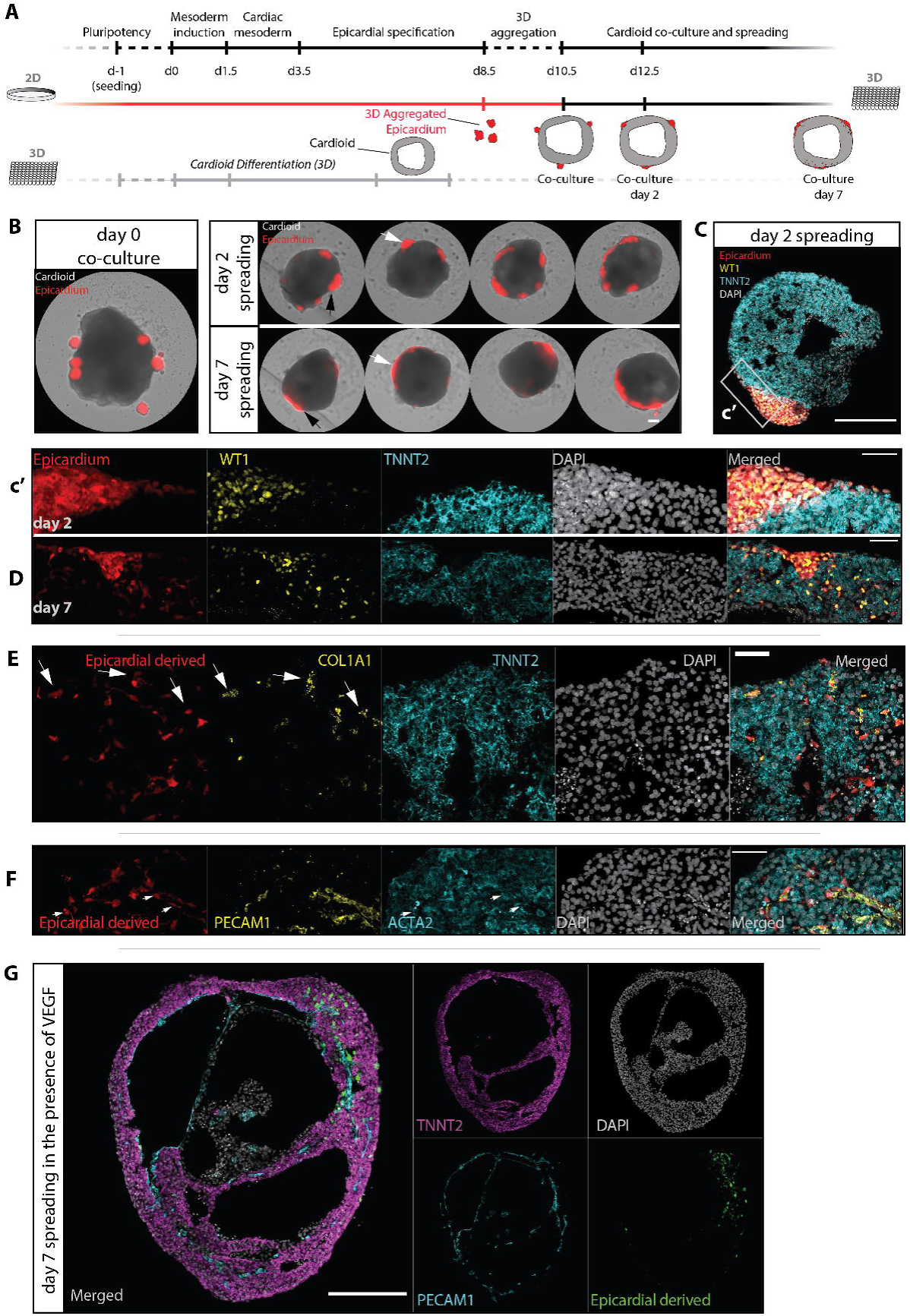
Epicardium Interacts with Cardioids and Migrates into the CM Layer. **(A-B)** Developmentally aligned protocol schematic showing fluorescently labeled 3D aggregates of 2D epicardial differentiations co-cultured with time matched cardioids. Arrows indicate morphology changes and spreading over time. Scale bar: 200 μm. **(C-D)** Confocal images of TNNT2 (CMs) and WT1 (epicardium) of 2- and 7-day co-cultures of epicardium and cardioids. Scale bars: 200 μm (C) and 50 μm (c’ and D). **(E-F)** Confocal images of COL1A1+ (E) and ACTA2+ (F) epicardial derivatives after 7 days of co-culture with cardioids containing ECs (PECAM1+). Scale bars: 50 μm. **(G)** Confocal images of epicardial derivatives after 7 days of co-culture with cardioids (TNNT2+) encompassing EC (PECAM1+) inner lining in the presence of 100 ng/ml VEGF-A. Scale bar: 200 μm.

To model the process of epicardial engulfment of the myocardium, we co-cultured cardioids with epicardial aggregates. We used basic media conditions without exogenous TGF-b, FGF and PDGF to test whether endogenous expression of these signaling factors by cardioids (Figure 7A,B, Figure S7F) is sufficient to stimulate epicardial interaction. We observed spreading of epicardial cells on top of cardioids within 2-7 days (Figure 7B-D, Figure S8C). After 7 days and without additional growth factors, epicardial cells interacted with the CM layer, migrated into it, and differentiated (Figure 7E-G, Figure S8C-F). The migrating cells downregulated epicardial WT1 and upregulated the SMC and CF markers ACTA2 (Figure 7F, Figure S8E) and COL1A1 (Figure 7F, Figure S8D,F), as occurs *in vivo* (Cao et al., 2019; Simões and Riley, 2018). Strikingly, some of these migrating epicardium-derived cells started to interact with cardioid ECs (Figure 7F,G, Figure S8E,F). We concluded that co-culture in the absence of external signals triggers intrinsic spreading of epicardium on cardioids, its inward migration, differentiation and interactions with CMs and ECs.

In conclusion, we have established a high-throughput human cardioid platform with intrinsic self-organization into patterned layers and 3D structures reminiscent of an early human heart chamber. We also show that our platform can be used to model cardiac cavity defects and study separate cellular processes and mechanisms underlying development of the three major cardiac lineages in the human heart.

## DISCUSSION

The variability and complexity of self-organizing organoid systems still hinders quantitative modelling of morphogenetic defects (Little and Combes, 2019). In cardioids, we address this challenge by omitting exogenous ECM and using a high-throughput approach to reach optimal conditions (Supplemental Videos 7, 8). We further increased reproducibility by tightly controlling the stepwise incorporation of the three main cardiac lineages into cardioids. This approach allows dissecting, with high statistical power, when and where the functions of specific factors are required. The simplicity of the system that can contain either one, two, or three cardiac lineages, without interference of non-cardiac tissues, makes it possible to strip self-organization and its underlying molecular and cell biological mechanisms to its bare essentials. Complexity in cardioids can therefore be tailored to the biological question asked. This is an important advantage for an organoid model as complex biological systems often employ redundant mechanisms that are otherwise challenging to tease apart.

Cardioids, as all other self-organizing organoid systems, recapitulate some aspects of development but also differ from embryogenesis in others. Self-organization encompasses only a subset of intrinsic developmental mechanisms, which are sufficient to recapitulate aspects of the *in vivo*-like architecture (Sasai, 2013). Consequently, using cardioids, we showed that cardiac mesoderm alone is sufficient to form a chamber-like cavity *in vitro*. We propose that this cavity could be analogous to the cavity of the heart tube and early heart chamber. *In vivo*, the first cavity arises from foregut endoderm-assisted migration and fusion of bilateral cardiac mesoderm and endocardial tubes into a single heart tube (Abu-Issa and Kirby, 2007). However, bilateral heart tubes and chambers can form in the absence of either endocardium (Ferdous et al., 2009) or foregut endoderm constriction (DeHaan and DeHaan, 1959; Li et al., 2004), but the mechanism is still unknown. This indicates the inherent capability of cardiac mesoderm to intrinsically form cavities and chambers *in vivo* (Ivanovitch et al., 2017), which is in agreement with the self-organization we observed in cardioids and chick embryo explants *in vitro*. Lateral plate mesoderm, a subtype closely related to cardiac mesoderm, has a similar potential to form a cavity − the pericardial body cavity (Schlueter and Mikawa, 2018). Thus, cavitation is a more general characteristic of mesoderm that could be called upon in embryos with a foregut defect. Finally, the HAND1 KO cavity phenotype in cardioids is consistent with the hypoplastic left ventricular chamber phenotype in Hand1 KO mice (McFadden, 2004; Riley et al., 1998; Risebro et al., 2006), and with the Hypoplastic Left Heart Syndrome chamber cavity phenotype in humans (Grossfeld et al., 2019), demonstrating the modelling potential of cardioids.

We used the cardioid platform to demonstrate that WNT, BMP and VEGF drive their self-organization. These pathways are known to regulate cardiac specification *in vivo* and *in vitro* (Meilhac and Buckingham, 2018), but whether and at what stage they control cardiac patterning and morphogenesis was unclear. The surprising finding that early mesodermal WNT controls later cardiac self-organization is consistent with early cardiac lineage diversification during mesoderm induction *in vivo* (Garcia-Martinez and Schoenwolf, 1993; Lescroart et al., 2018). Patterning and morphogenesis occur in parallel with specification, but they are not necessarily linked. In agreement with this notion, cavities can self-organize in the absence of cardiac specification and in HAND1 KO cardioids there is a defect in self-organization but not in CM specification. Conversely, inhibition of WNT signaling at the cardiac mesoderm stage is essential for CM specification but does not regulate cardioid self-organization. Cardioids are therefore a powerful system to intrinsically dissect regulation of specification and morphogenesis. At the same time, cardioids are simple enough to determine sufficiency of a factor for one of these processes and are thus complementary to more complex systems.

We found that WNT and VEGF control CM and EC self-organization in cardioids. *In vivo*, cardiac ECs first form endocardial tubes, later become separated from the outer CM tube by an ECM-filled (cardiac jelly) interspace, and finally form the inner lining of the heart chambers (Abu-Issa and Kirby, 2007; Ivanovitch et al., 2017). How signaling coordinates these patterns and morphogenetic processes with specification was unclear. In cardioids, the patterning and morphogenesis of CM and EC lineages is controlled by the dosage of WNT at the earliest stage of mesodermal differentiation, and by VEGF that directs both specification and patterning of the EC layer in cardiac mesoderm. When ECs and CMs are aggregated in microtissues, they do not form separate layers and lining (Giacomelli et al., 2017). Generation of a separate EC layer and lining is crucial for activation of mechanosensing in the context of a chamber *in vivo*. Cardiac chamber mechanobiology is required for physiological EC and CM crosstalk, driving the next stages of heart development like trabeculation, myocardial compaction and interaction with epicardium (Wilsbacher and McNally, 2016). Cardioids are therefore a promising system to study the underlying mechanisms of CM and EC patterning and crosstalk in the context of a beating chamber.

During development, the (pro-)epicardium makes contact with the early heart chambers, engulfs them, and concomitantly differentiates and migrates into the myocardium (Cao et al., 2019; Simões and Riley, 2018). By co-culturing cardioids with epicardium, we observed epicardial spreading, migration and differentiation reminiscent of these processes *in vivo*. Epicardial and CM co-cultures have been studied before using microtissues (Guadix et al., 2017), but not in the context of a cardiac chamber-like model. This aspect is crucial because the crosstalk between derivatives of the epicardial, EC and CM lineages is dependent on the mechanobiology of the heart chamber (Wilsbacher and McNally, 2016). We therefore propose that self-organization of the three cardiac lineages in chamber-like cardioids will be important to reignite the physiological crosstalk that drives growth and maturation of the heart as *in vivo*.

Overall, cardioids provide a unique foundation for the incorporation of additional cardiac lineages by signaling controlled self-organization (Kelly et al., 2014). We postulate that increased complexity using developmental principles *in vitro* will induce further maturation and functionalization providing models that could transform cardiovascular research. The cardioid platform has a wide potential to explore fundamental mechanisms of self-organization and congenital defects, as well as to generate future mature and complex human heart models suitable for drug discovery and regenerative medicine.

## ACKNOWLEDGEMENTS

We thank all laboratory members for help and discussions and in particular Katarzyna Warczok for lab management. We are grateful to the VBC, IMP/IMBA Core and IMBA SCCF facilities for their services, the Allen Institute for cell lines, the Knoblich lab and Joshua A. Bagley for cell lines, reagents and discussions, and Reiner Wimmer and Josef Penninger for sharing data and reagents. We are thankful to Anna Kicheva and the Kicheva lab for providing training on chicken embryo work. We would also like to thank the Tanaka and Gerlich labs for sharing reagents. We are grateful to Paulina Latos, Alex Stark, Luisa Cochella, Juergen Knoblich and Elly Tanaka for critical reading and comments on the manuscript. This work was funded by the Austrian Academy of Sciences.

## AUTHOR CONTRIBUTIONS

P.H., S.J., N.P. and S.M. co-designed experiments and co-wrote the paper. P.H. optimized CM and EC differentiations in 2D, co-differentiations in 3D aggregates and in cardioids, and set up the high-throughput analysis pipeline; S.J. made the initial observation of cardiac cavity formation and optimized CM cardioid generation; and N.P. developed the epicardial differentiations in 2D, 3D and in cardioids. M.N. helped with the bioinformatic analysis and all other authors performed experiments. S.M. supervised the study.

## METHODS

Details on hPSC differentiation into cardiomyocytes in 2D and 3D aggregates, generation of cardioids, generation of cardioids with endothelial lining, cell-line-dependent CHIR99021 concentrations, epicardial co-culture with cardioids, and 2D anterior endothelial cell differentiation will be uploaded in a separate BioRxiv submission to allow for more detailed description of the protocols.

### General human pluripotent stem cell culture

Human pluripotent stem cell lines (WT H9, WiCell and constitutively fluorescent H9 clones (Wimmer et al., 2019) WT and modified WTC, Allen Institute for Cell Science) were cultured in a modified in-house medium based on the E8 culture system (Chen et al., 2011). Cells were grown on either Corning or Eppendorf tissue culture-treated plates coated with Vitronectin XF (Stem Cell Technologies #7180) and passaged using either TrypLE Express Enzyme (Gibco, #12605010) or PBS-EDTA (Biological Industries, 01-862-1B) every 2-4 days at ∼70% confluency. Cells were routinely tested for Mycoplasma.

### ECM molecules used

Vitronectin (10 µg/ml), Laminin-511 E8 fragment (Takara Bio, #T303, 0.05 – 2 µg/ml) and Laminin-521 (Biolaminin, # LN521-02, 0.1 – 5 µg/ml) were either used to pre-coat wells or added to the cell suspension prior to seeding. Further cardiomyocyte differentiation was performed as described above.

### Culture of Human Cardiac Microvascular Endothelial Cells

Human cardiac microvascular endothelial cells (HCMEC) were obtained from PromoCell (PC-C-12285 HCMEC-c) and cultured according to manufacturer’s instructions using Endothelial Cell Growth Medium MV (PromoCell, #PC-C-22020). For Smart-Seq2 analysis, HCMEC were dissociated with TrypLE Express Enzyme and FACS-sorted into home-made lysis buffer.

### Chick cardiac mesoderm explant culture

Explants from the cardiogenic region of developing chicken (*Gallus gallus*) embryos were isolated at Hamburger and Hamilton stage 7-8 and cultured for 24h at 37°C in cardiac mesoderm (BFIIWPRa) media. Subsequently explants were embedded, cryosectioned and immunostained for further analyses as described below.

### Cryosectioning

Cryosectioning was done based on (Bagley et al., 2017). Briefly, 4% PFA-fixed tissues were cryoprotected with 30% sucrose in PBS overnight at 4°C and embedded the next day using O.C.T. cryoembedding medium (Scigen, #4586K1). Embedded tissues were frozen using a metal surface submerged in liquid nitrogen and tissues were stored in a −80°C freezer until sectioning on a Leica cryostat. Sections were collected on Ultra Plus slides and kept at −20°C or − 80°C until immunostaining. O.C.T. was removed by washing with PBS before continuing with the immunostaining protocol.

### Immunostaining

Following fixation with 4% PFA (Sigma-Aldrich, #16005) specimens were washed twice in 1x PBS and the 3D constructs additionally once in PBS/Tween20 (0.1%, Sigma-Aldrich, #P1379) for at least 15 min each. Tissues were incubated in blocking solution consisting of PBS (Gibco, #14190094) with 4% goat (Bio-Rad Laboratories, #C07SA) or donkey serum (Bio-Rad Laboratories, #C06SB) and 0.2% Triton X-100 (Sigma-Aldrich, #T8787) for at least 15 minutes. The primary antibody was subsequently applied in above blocking buffer for 1-3 hours at room temperature or overnight at 4°C in the case of the 2D samples and 2 days at 4°C on a shaker for 3D samples. Following washing twice with PBS/Tween20, 3D tissues were incubated with the secondary antibody solution at 4°C on a shaker for 2 more days while 2D samples were incubated up to 2 hours at room temperature. Following these washing steps and an additional PBS wash, tissues were ready for analysis or storage at 4°C in PBS, while slides were mounted using fluorescence mounting medium (Dako Agilent Pathology Solutions, #S3023). 3D tissues were cleared with FocusClear (CellExplorer Labs, #FC-101) prior to imaging.

### Dextran and Fluo-4 incorporation assays

A 4.4kDa Dextran conjugated with TAMRA (Sigma Aldrich, T1037) was added to the cardioid culture between 64h and 90h. Cardioids were live imaged as cavities started to form. To image and analyze calcium-transients, 3D cardioids and 2D CM were loaded with Fluo-4 AM (Thermo Fisher Scientific, #F14217). After a 15-min incubation, cardioids were incubated for another 15 min in Tyrode’s salt solution (Sigma Aldrich, #T2397). Subsequently, cardioids were imaged live and videos were analyzed with FIJI software (Schindelin et al., 2012) to acquire the signal intensity (F) of regions of interest and background (F0).

### Contraction characteristics measurements

2D CM and 3D cardioids were live-imaged and videos were analyzed with a published algorithm (Huebsch et al., 2014) to determine contraction velocity and beating rate.

### Image acquisition and analysis

Fixed whole mounts and sections were imaged with point scanning (upright Zeiss LSM800 Axio Imager with an Apochromat 20x objective lens at 1x magnification) and spinning disk confocal microscopes (Olympus spinning disk system based on a IX3 Series (IX83) inverted microscope, equipped with a Yokogawa W1 spinning disc), or a widefield microscope (Zeiss Axio Imager 2, Axio Vert A1). Live imaging experiments were performed using a Zeiss Celldiscoverer 7 or above-mentioned spinning disk microscope. For high-throughput imaging and analysis, images were taken with a Celigo Imaging Cytometer microscope (Nexcelom Biosciences, LLC) and analyzed with custom-made scripts written for the Fiji software (Schindelin et al., 2012).

### Statistics

Data is presented as Mean +/- SD. To calculate significant differences, data was analyzed for normality and lognormality using the D’Agostino & Pearson and the Shapiro-Wilk test in Prism 8 software (GraphPad Software Inc.). If normally distributed, a parametric test (two-tailed t-test, one-way ANOVA) was performed to determine significant differences. If not normally distributed, a non-parametric test (two-tailed Mann-Whitney, Kruskal-Wallis) was performed using Prism 8 software. Corrections for multiple comparisons using statistical hypothesis testing (Tukey test for parametric and Dunn’s test for non-parametric) was performed using Prism 8 software. The p-values for significant differences are visualized as: *: p<0.05, **: p<0.01, ***: p<0.001, ****: p<0.0001.

### Flow cytometry

Cells were dissociated using the CM dissociation kit (Stem Cell Technologies, #05025). After centrifugation for 3 min at 130g, cells were resuspended in 300 µl PBS supplemented with 0.5 mM EDTA (Biological Industries, #01-862-1B) and 10% FBS (PAA Laboratories, #A15-108). Cells were acquired with a FACS LSR Fortessa II (BD) and analysed with FlowJo V10 (FlowJo, LLC) software. FACS sorting was performed using a Sony SH800 Cell Sorter (Sony Biotechnology).

### RNA isolation and RNA-seq/Smart-Seq2 preparation

RNA was isolated with the RNeasy Mini Kit (Qiagen, #74104). Generation of the RNA-seq libraries was performed according to the manufacturer’s instructions with QuantSeq 3’ mRNA-Seq Library Prep Kit FWD (Lexogen GmbH, #015). After the preparation of the libraries, samples were checked for an adequate size distribution with a fragment analyzer (Advanced Analytical Technologies, Inc) and were submitted to the Vienna Biocenter Core Facilities (VBCF) Next-Generation-Sequencing (NGS) facility for sequencing. For Smart-Seq2 analysis, 400 cells were sorted into lysis buffer and stored at −80°C until further processing. Samples were QC’d/libraries prepared and sequenced by the VBCF NGS facility using a home-made Smart-Seq2 kit.

### Bioinformatic analysis

Trimming was performed for Smart-Seq2 experiments using trim-galore v0.5.0 and for QuantSeq 3’ mRNA-Seq experiments using BBDuk v38.06 (ref=polyA.fa.gz,truseq.fa.gz k=13 ktrim=r useshortkmers=t mink=5 qtrim=r trimq=10 minlength=20). Reads mapping to abundant sequences included in the iGenomes UCSC hg38 reference (human rDNA, human mitochondrial chromosome, phiX174 genome, adapter) were removed using bowtie2 v2.3.4.1 alignment. Remaining reads were analyzed using genome and UCSC gene annotation provided in the iGenomes UCSC hg38 bundle (https://support.illumina.com/sequencing/sequencing_software/igenome.html). Reads were aligned to the hg38 genome using star v2.6.0c and reads in genes were counted with featureCounts (subread v1.6.2) using strand-specific read counting for QuantSeq experiments (-s 1). Differential gene expression analysis on raw counts, and principal component analysis on variance-stabilized, transformed count data were performed using DESeq2 v1.18.1. Functional annotation enrichment analysis of differentially expressed genes was conducted using clusterprofiler v3.6.0 in R v3.4.1. Bulk tissue cell type deconvolution was performed using MuSiC v0.1.1 (Wang et al., 2019). We used cell type-specific marker genes and a cell-type specific single-cell expression reference for the human developing heart (Cui et al., 2019). Bulk cardioid RNA-seq samples were processed with the previously described pipeline using the hg19 UCSC iGenomes reference to match the published data. The proportion of cell types from developing heart in bulk cardioid RNA-seq samples was estimated and MuSiC estimated proportions were visualized in a heatmap.

### Proteomics

Cells were lysed in 8M urea 100mM 4-(2-hydroxyethyl)-1-piperazineethanesulfonic acid buffer (HEPES), reduced with 10 mM 1,4-dithioerythritol with 1U Benzonase (Merck KGaA, #1.01654.0001) and alkylated with 20 mM 2-iodoacetamide. Digests were carried out in 4M urea 100mM HEPES with LysC (Wako, #121-05063, 1/100 (w/w) protease/substrates) for 3 h at 37C and subsequent trypsin digest (Promega, #V5280, 1/100 (w/w) protease/substrates) overnight at 37C. Peptides were desalted using reverse-phase solid phase extraction cartridges (Sep-Pak C-18, Waters, #186000308), dried under vacuum, reconstituted in HEPES to a neutral pH and labeled with TMT10-plex (Thermo Fisher Scientific, #90110) according to manufacturer’s instructions. TMT labeled peptides were pooled in equal amounts and fractionated by high pH reversed phase chromatography (UPLC Peptide CSH C18 column, 130Å, 1.7 µm, 1 mm x 150 mm, ACQUITY) to obtain 10 final fractions.

The samples were separated by reversed phase chromatography (75 um × 250 mm PepMap C18, particle size 5 um, Thermo Fisher Scientific), developing a linear gradient from 2% to 80% acetonitrile in 0.1% formic acid within 60 minutes (RSLC nano, Dionex – Thermo Fisher Scientific) and analyzed by MS/MS, using electrospray ionization tandem mass spectrometry (Orbitrap QExactive HFX, Thermo Fisher Scientific). The instrument was operated with the following parameters: MS1 resolution 120,000; MS1 AGC target 3e6; MS1 maximum inject time 50ms; MS1 scan range 380 to 1650 m/z; MS2 resolution 45,000; MS2 AGC target 1e5; Maximum inject time 250; TopN 10; Isolation window 0.7 m/z; Fixed first mass 110 m/z; Normalized collision energy 35; Minimum AGC target 1e4; Peptide match preferred; Exclude isotope on; Dynamic exclusion 30s.

All MS/MS data were processed and analysed using Proteome Discoverer 2.3 (PD 2.3.0.484, Thermo Scientific), searched using MSAmanda v2.0.0.14114 (Dorfer et al., 2014) against the Homo sapiens database (SwissProt TaxID=9606) (v2017-10-25). Maximal missed cleavages: 2, with iodoacetamide derivative on cysteine and peptide N-terminal ten-plex tandem mass tag (fixed mod.); oxidation on methionine, ten-plex tandem mass tag on lysine (variable mod.). Peptide mass tolerance: ±5 ppm; fragment mass tolerance: ±15 ppm. Filtered to 1% FDR on protein and peptide level using Percolator; reporter ions were quantified using IMP Hyperplex (Doblmann et al., 2019) (https://ms.imp.ac.at/index.php?action=hyperplex).

### Generation of MYL7-GFP/CDH5-Tomato double reporter line

The endogenously tagged WTC MYL7-GFP hPSC line was obtained from the Allen Institute for Cell Science (Cell Line ID: AICS-0052). A gBlock of the CDH5 promoter sequence (−1135 to − 5 relative to TSS) (Prandini et al., 2005) was ordered from Integrated DNA Technologies, Inc. and cloned according to Bagley et al., into a modified backbone of a vector that integrates into the AAVS1 locus with TALEN technology (Hockemeyer et al., 2009). The modified backbone contained flanking tandem repeats of the core chicken HS4 insulator (2xCHS4). Thus, the following reporter expression cassette was inserted in the AAVS1 locus: 2xCHS4-CDH5promoter-dTomato-WPRE-SV40-2xCHS4. Nucleofection and clone picking/validation was done as in (Bagley et al., 2017).

### Generation of HAND1 and NKX2-5 Knock Out Cell Lines

HAND1 and NKX2.5 were knocked out in H9 cells using CRISPR/Cas9. sgRNAs for target sites were identified using the Sanger Institute Genome Editing (WGE) website, as well as the Benchling sgRNA designing tool. (HAND1_sgRNA1: GAGCATTAACAGCGCATTCG; NKX2.5_sgRNA1: GACGCACACTTGGCCGGTGA; NKX2.5_sgRNA2: ACTTGGCCGGTGAAGGCGCG). sgRNAs were cloned into pSpCas9(BB)-2A-Puro (PX459) V2.0 (Feng Zhang Lab; Addgene plasmid #62988; http://n2t.net/addgene:62988; RRID:Addgene_62988) according to the Zhang Lab General Cloning Protocol (Ran et al., 2013). Cells were transfected using the P3 Primary Cell 4D-Nucleofector™ X Kit S (Lonza-BioResearch, Cat #: V4XP-3032) and Amaxa™ 4D-Nucleofector™ (Lonza-BioResearch). Post nucleofection, cells were incubated in E8 supplemented with 10μM Y-27632 (Cat #72302) for 24h. After that period, cells were selected with puromycin (concentration 0.2ng/μL; (Sigma-Aldrich, Cat #P8833) for 48h. Following this treatment, the cell culture media was changed back to E8 supplemented with 10μM Y-27632 (Cat #72302) to promote re-growth. Once the cells formed colonies, they were picked and transferred into a 96w-plate (Corning, Cat #CLS3370). Successful editing was first assessed on a pool level. Subsequently, single colonies were genotyped two times independently in order to be confirmed a successful knock out. Genome editing on a pool and clonal level was assessed by Synthego’s online tool ICE (https://ice.synthego.com/#/).

Primers:

HAND1_G1_forward 5’-CACCGAGCATTAACAGCGCATTCG-3’

HAND1_G1_reverse 5’-AAACCGAATGCGCTGTTAATGCTCC-3’

NKX2.5_G1_forward 5’-CACCGGACGCACACTTGGCCGGTGA-3’

NKX2.5_G1_reverse 5’-AAACTCACCGGCCAAGTGTGCGTC C-3’

NKX2.5_G2_forward 5’-CACCGACTTGGCCGGTGAAGGCGCG-3’

NKX2.5_G2_reverse 5’-AAACCGCGCCTTCACCGGCCAAGT-3’

### Antibodies

The following antibodies were used: TNNT2 (Thermo Scientific, MS-295-P), VE-Cadherin (CDH5) (Cell Signaling Technology, 2500S), PECAM1/CD31 (Agilent Technologies, M082329-2), PECAM1/CD31 (R&D Systems, AF806-SP), HAND1 (R&D Systems, AF3168), Cleaved Caspase-3 (Cell Signaling Technology, 9661s), WT1 (Abcam, ab89901, KI67 (BD Biosciences, 556003), Phalloidin (Invitrogen, A12380), N-Cadherin (BD Biosciences, BD610920), Fibronectin (Sigma Aldrich, F3648), SOX18 (Novus Biologicals, NBP2-58004), VIM (DSHB at the University lowa, AMF-17b), ACTA2 (α-SMA) (Sigma-Aldrich, A2547), Collagen Type I (DSHB at the University lowa, SP1-D8 and Invitrogen, PA529569), EOMES (R&D Systems, AF6166-SP), SOX17 (R&D Systems, AF1924-SP), Beta-Catenin (DSHB at the University lowa, PY489-B-Catenin), SOX2 (R&D Systems, RD-AF2018), TBX5 (Sigma-Aldrich, HPA008786).

### Data availability statement

Patsch et al., HUVEC and Vascular Organoid EC Smart-seq2 data was taken from (Wimmer et al., 2019) for analysis. RNA-seq data has been deposited to NCBI Gene Expression Omnibus and is accessible through the GEO accession number GSE148025 (https://www.ncbi.nlm.nih.gov/geo/query/acc.cgi?acc=GSE148025). The mass spectrometry proteomics data have been deposited to the ProteomeXchange Consortium via the PRIDE (Perez-Riverol et al., 2019) partner repository with the dataset identifier PXD018306. Other data is available upon request.

---

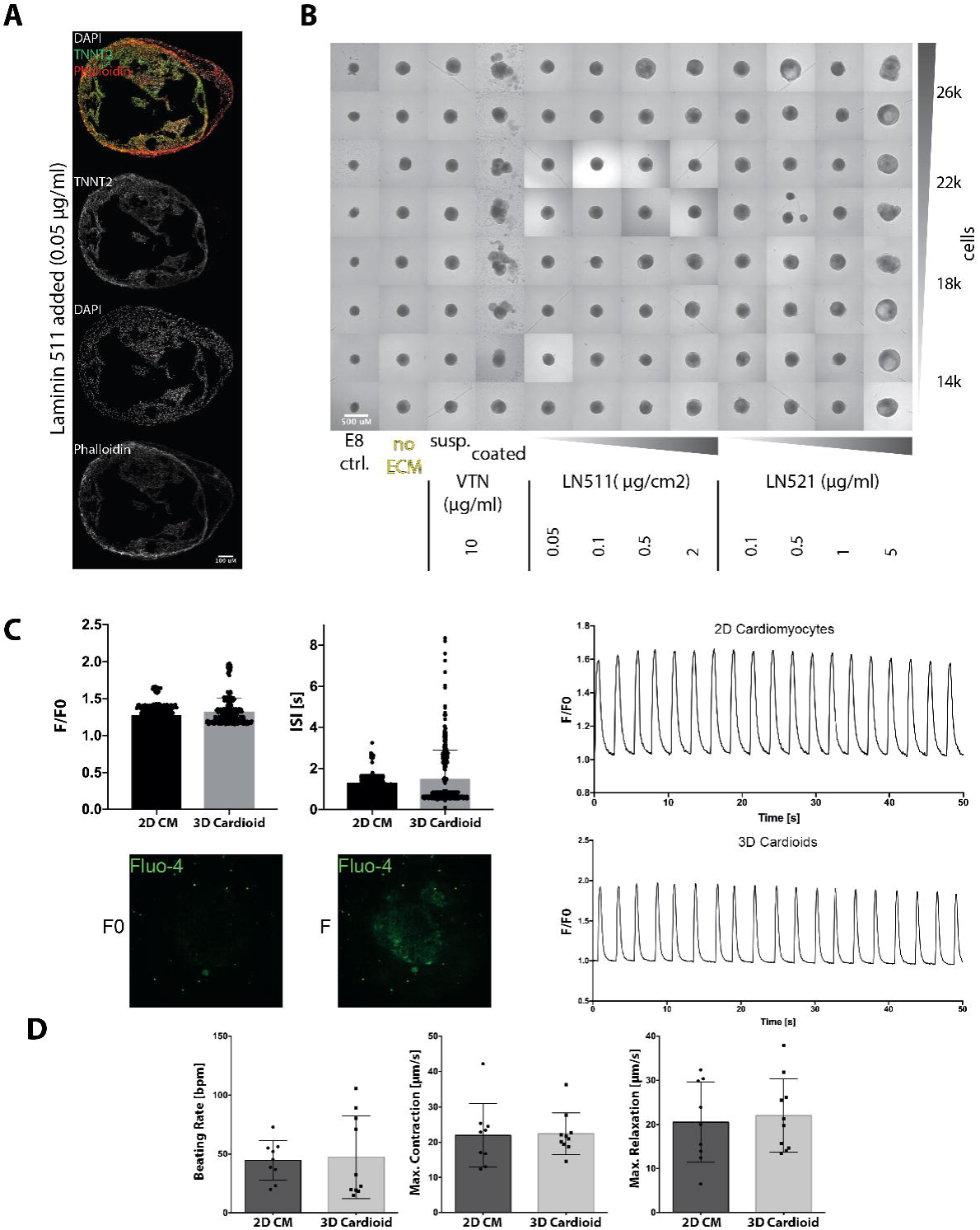

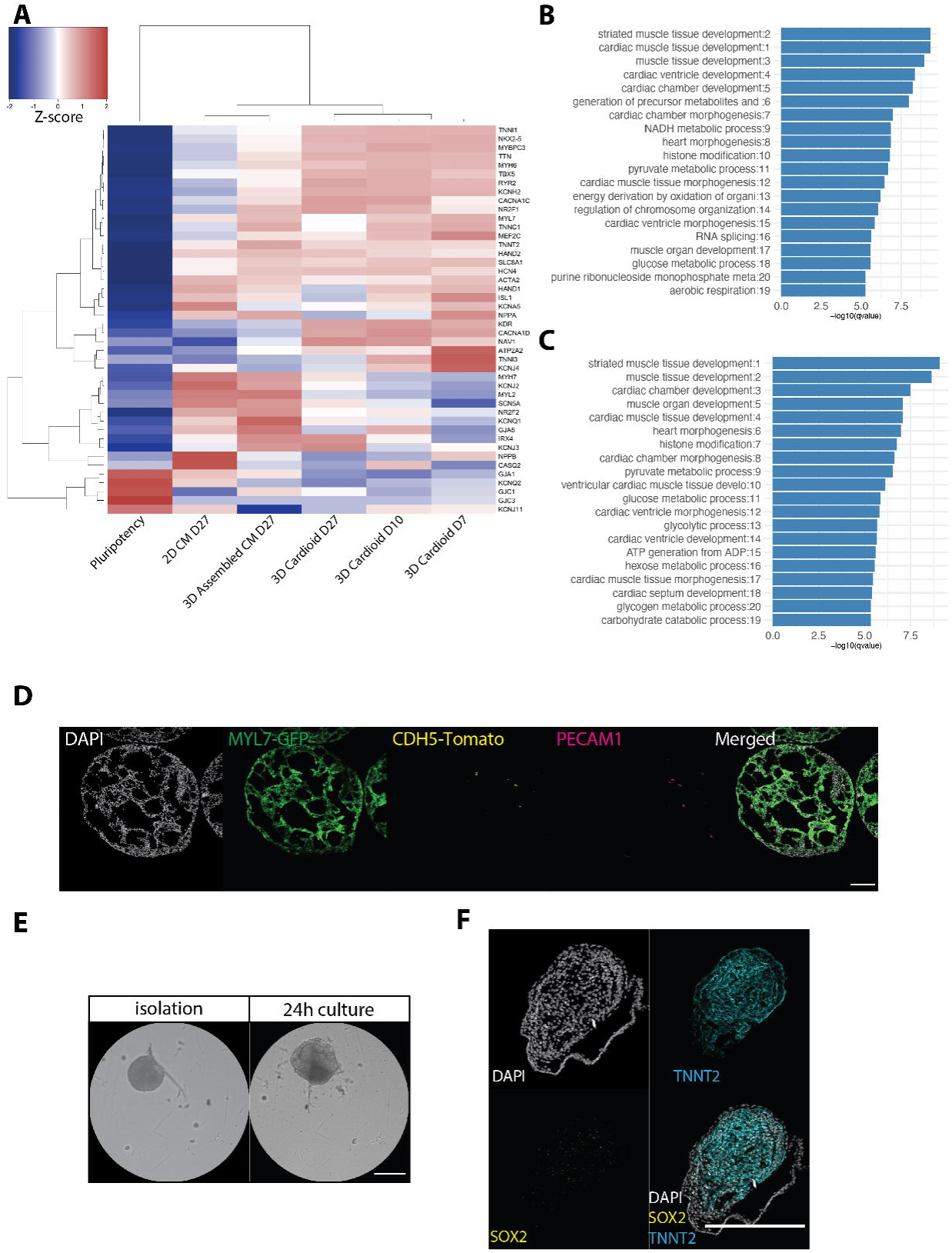

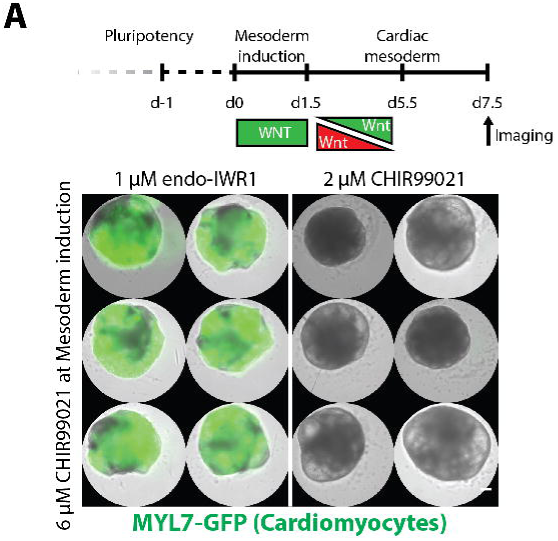

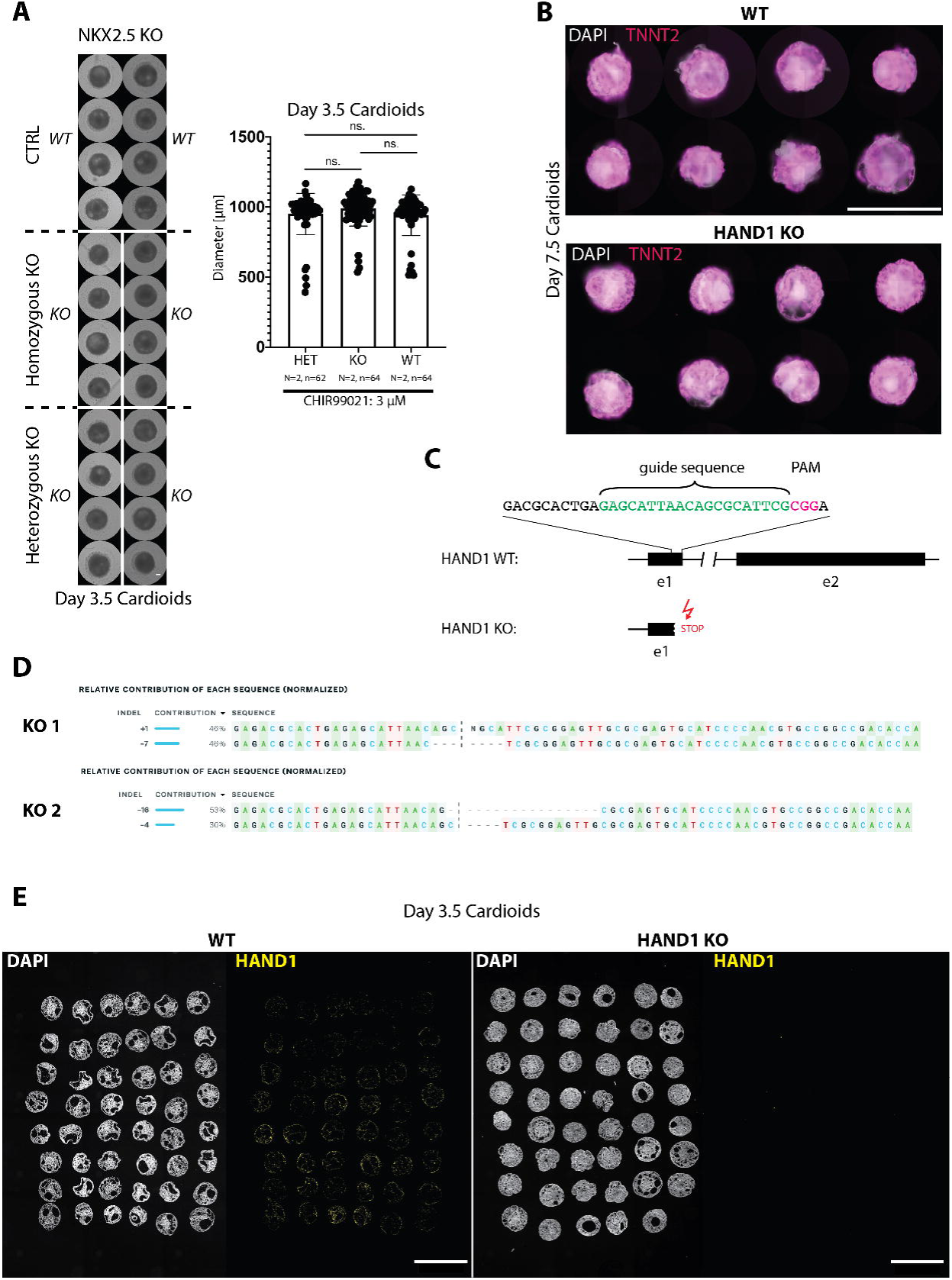

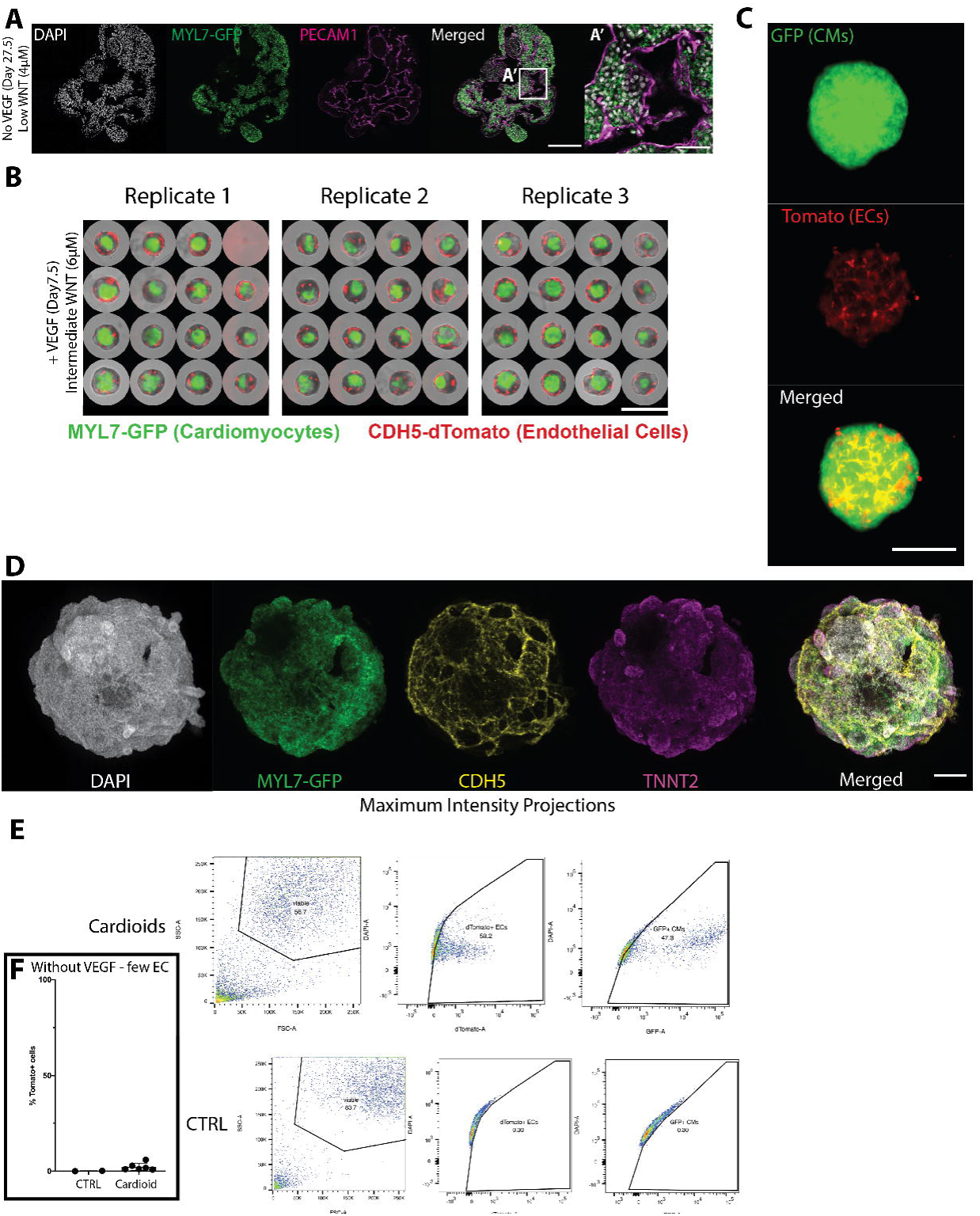

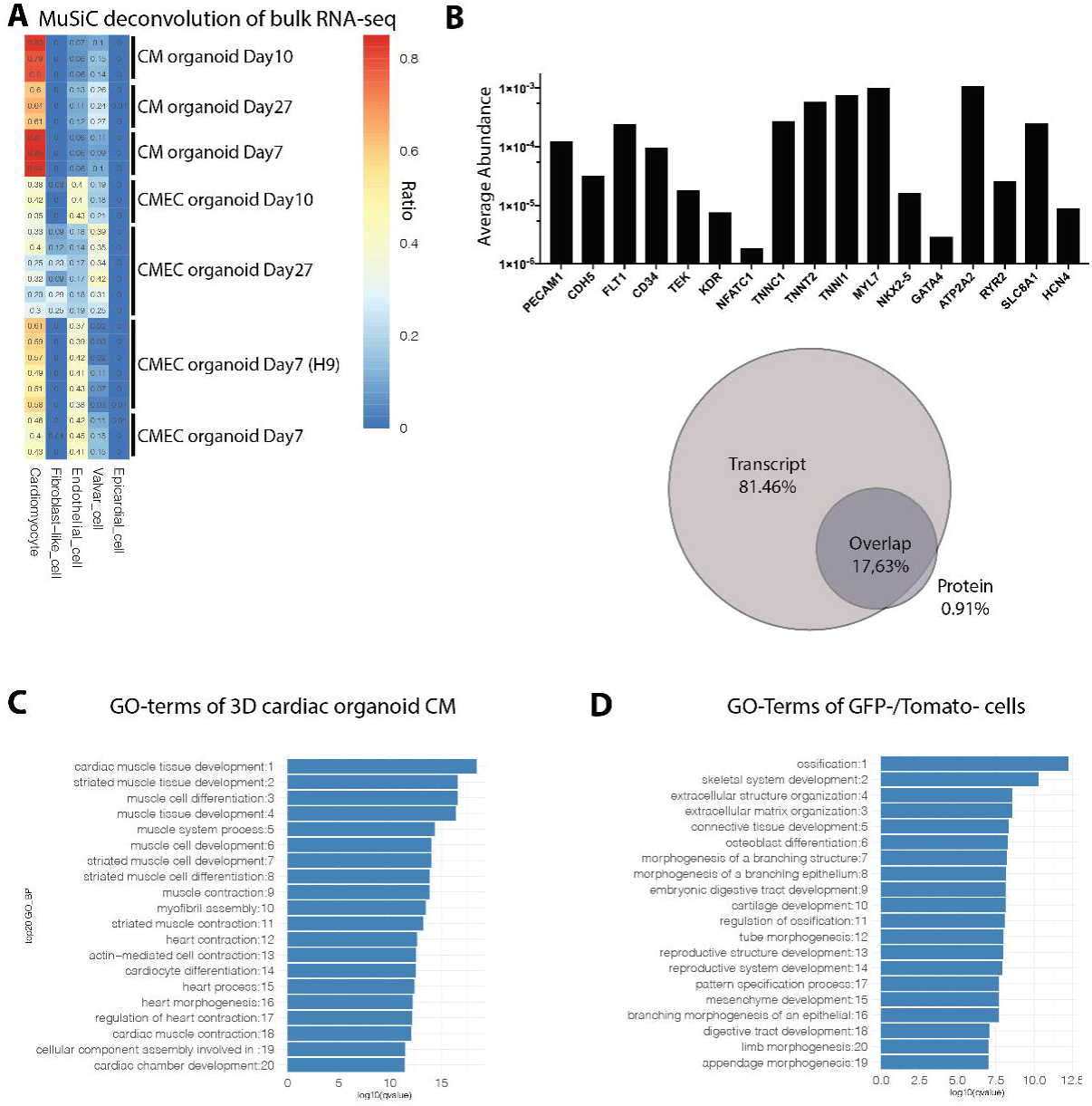

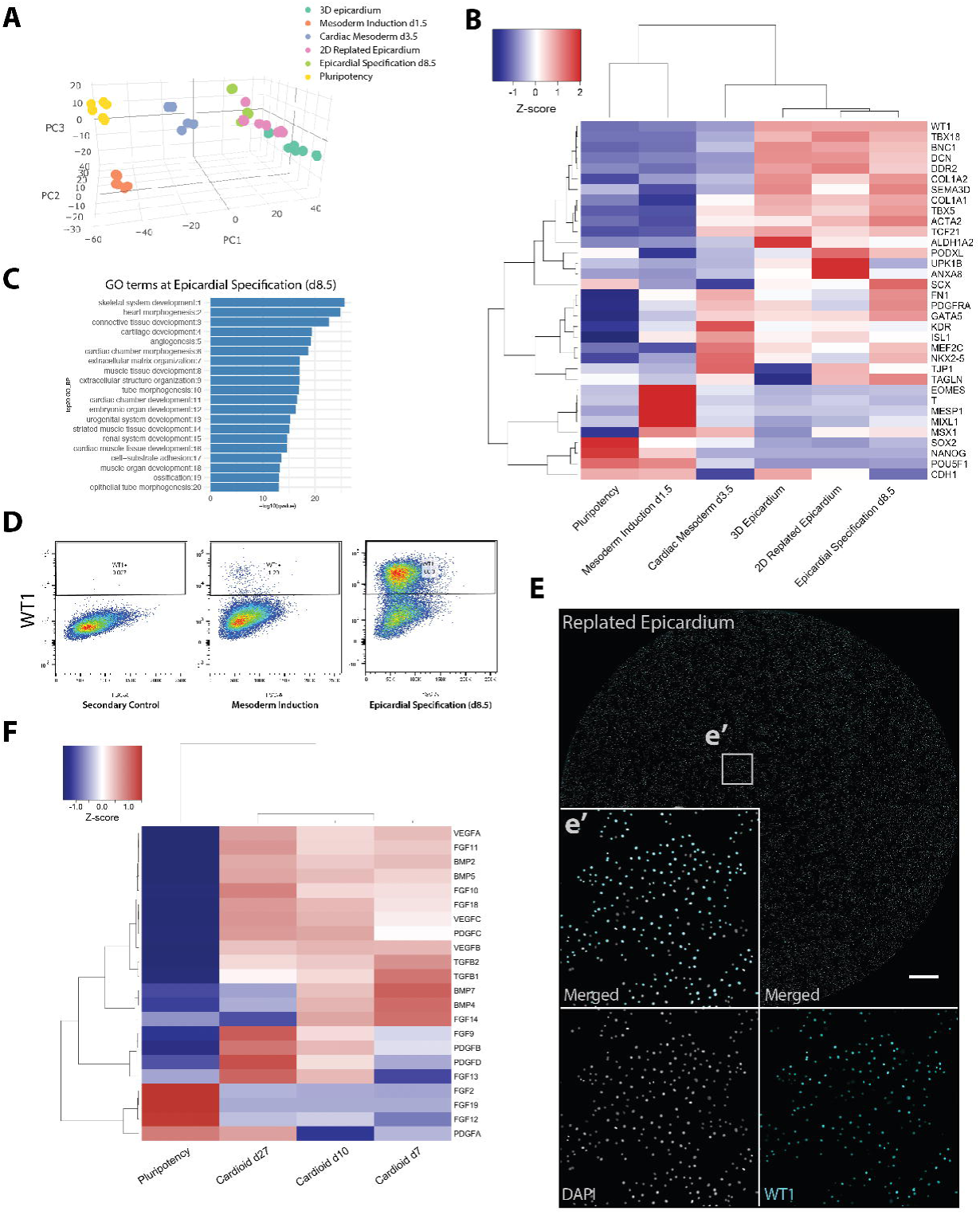

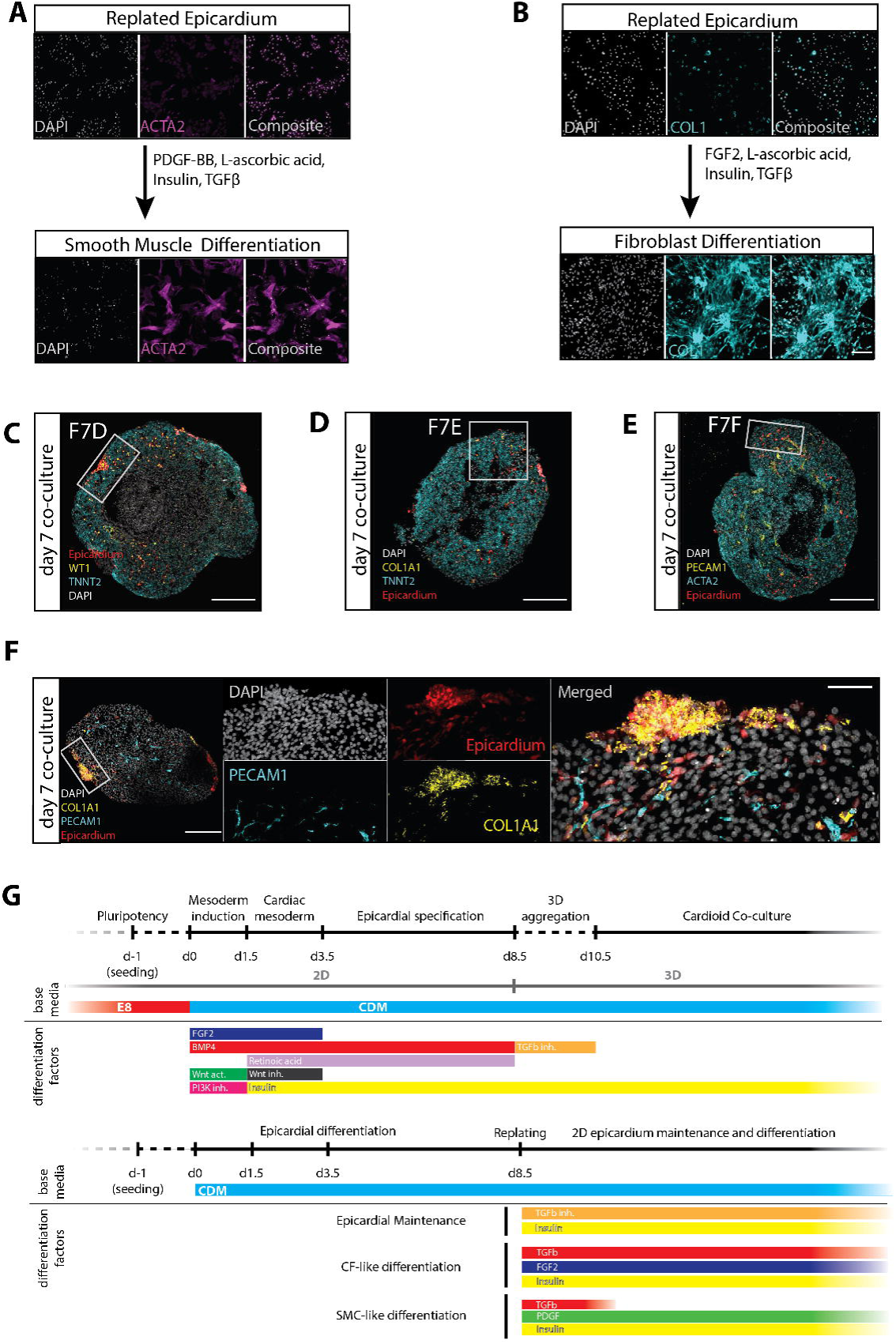

