## Supplemental Figure Legends for "Cardioids reveal self-organizing principles of human cardiogenesis"

**Figure S1. Generation and Functional Characterization of Chamber-like Structures. Related to Figure 1. (A)** Example of a beating chamber-like structure (day 8.5) containing a large cavity derived by adding laminin LN511 (0.1 µg/ml) to the cell suspension during seeding prior to differentiation in a 2D flat bottom well. Scalebar: 100µm. **(B)** When cardiomyocyte differentiation is performed in ultra-low attachment plates, beating 3D structures with cavities form in the absence of extrinsic ECM molecules. Timepoint: day 8.5. Scalebar: 500µm. **(C)** Chamber-like structures and 2D cardiomyocytes (CM) display calcium transients that are similar. Calcium transients analyzed by loading the cells with Fluo-4-AM and analyzing the fluorescence intensity over time. F/F0: Fluorescence intensity over background. ISI: inter-spike-interval. **(D)** Analysis of beating parameters using a published algorithm (Huebsch et al., 2015).

**Figure S2. Characterization of Cardioids and Chick Cardiac Mesoderm Explants. Related to Figure 2. (A) Gene** expression signatures of cardioids, aggregated cardiomyocytes and hPSCs at different timepoints. **(B)** GO terms upregulated in day 7 cardioids compared to day 7 CMs in 2D. **(C)** GO terms upregulated between day 7 in cardioids and day 27 3D aggregated CMs. **(D)** Day 7.5 cavity formation in cardioids sin the absence of endothelial cells (PECAM1+) upon treatment with Sunitinib (100 nM). Scalebar: 200 µm. **(E)** Chick cardiac mesoderm explants form a spherical structure with cavities *in vitro*. Scalebar: 200 µm. **(F)** Control chick cardiac mesoderm explants section (same CM TNNT+ structure as in Figure 2) shows near absence of the foregut marker SOX2. Scalebar: 200 µm.

**Figure S3. WNT Inhibition Controls Cardiac Mesoderm Specification but not Cavity Formation. Related to Figure 3. (A)** Wnt activation during the cardiac mesoderm stage inhibits CM-differentiation but not cavity expansion. Scalebar: 200 µm.

**Figure S4. NKX2-5 and HAND1 KO hPSC and Cardioid Characterization. Related to Figure 4. (A)** NKX2-5 KO cardioids do not show impaired cavity expansion. Scalebar: 200 µm. Quantification confirms no diameter reduction of NKX2-5 KO cardioids at day 3.5. **(B)** *HAND1* KO cardioids still express TNNT2 showing normal CM-differentiation capacity. Scalebar: 2500 µm. **(C)** Schematic of *HAND1* KO generation strategy. **(D)** Genotype of *HAND1* KO cell lines used for this study. Dotted line indicates the cutting site of CRISPR/Cas9-mediated editing. **(E)** Sections through 48 (WT) and 47 (KO) cardioids showing the increased number of cavities as well as HAND1 expression in WT cardioids only. Scalebars: 2000 µm

**Figure S5. Characterization of Cardioids with Endothelial Layers. Related to Figure 5. (A)** Cardioids generated with low WNT dosage (CHIR99021, 4µM) on day 27.5 showing inner EC lining the cavity and CMs. Scalebar: 200 µm. **(A')** Detail of inner EC cavity lining. Scalebar: 50 µm. **(B)** Three biological replicates showing reproducible generation of cardioids with separate CM and EC layers using an intermediate WNT dosage (CHIR99021, 6 µM). **(C)** ECs in aggregated CM microtissues build a rudimentary network within the cardiomyocytes, not around them. Scalebar: 400 µm. **(D)** EC build networks surrounding the cardioid. Max. Int. Projections. Scalebar: 200 µm. **(E)** Example FACS plots showing the distribution of cardiomyocytes (CM) and (EC) in co-differentiated cardioids at day 7.5 using the MYL7-GFP/CDH5-Tomato iPSC (WTC) line. CTRL: WT WTC line. **(F)** Without VEGF-A treatment, EC contribution is minimal.

**Figure S6. Characterization of Cardioid EC Identity. Related to Figure 6. (A)** MuSiC deconvolution (Wang et al., 2019) of bulk RNA-seq data of cardiac organoids showing CM/EC contributions similar to the FACS quantification and lack of ECs in CM-only cardioids.

Reference dataset: single cell RNA-seq of the developmental trajectory of the human heart (Cui et al., 2019). **(B)** Proteomic analysis of cardioids shows that almost all detected proteins are also detected in the transcriptome (day 7.5). Key markers of CMs and ECs are detected. **(C)** GO-terms of genes upregulated in CMs show terms related to cardiac muscle development. **(D)** GO-terms of genes upregulated in Non-CMs/Non-ECs show terms related to connective tissue development and ECM organization.

**Figure S7. Characterization of Epicardial Differentiation. Related to Figure 7. (A)** PCA of epicardium differentiation time course RNA-Seq. N>3 (H9 line) **(B)** Differential expression of pluripotency, mesoderm specification, cardiac mesoderm and epicardium specific genes, comparing day 8.5 (end of differentiation), 2D re-plated and 3D aggregated epicardium. **(C)** Top 20 Gene Ontology (GO) term enrichment of upregulated genes at the end of differentiation (day 8.5) compared to pluripotency. **(D)** FACS quantification of the WT1+ population at the end of differentiation (day 8.5) compared to mesoderm induction and secondary only control. **(E)** Whole well WT1 immunostaining of 2D re-plated epicardium. Scale bar: 1mm. e' is a 1x1 mm detail of E. **(F)** Differential expression of growth factors involved in epicardial sub-lineage specification in pluripotency and 7/10/27-day cardioids without co-culture.

**Figure S8. Epicardium Upregulates Fibroblast and Smooth muscle-like Markers *in vitro* after Differentiation. Related to Figure 7. (A-B)** ACTA2 (α-SMA) immunostaining of 2D re-plated epicardium and cells after SMC-like differentiation. **(B)** Type I Collagen immunostaining of 2D re-plated epicardium and cells after fibroblast differentiation. Scale bar: 200 μm. **(C-E)** Whole cardioid images of details from Figure 7D-F, Scale bars: 200 μm. **(F)** Confocal images of COL1A1+ epicardial-derivatives after 7 days of co-culture with cardioids containing COL1A1+ ECs (PECAM1+). Scale bars: 200 μm (overview) 50 μm (detail). **(G)**

Schematic of the epicardial differentiations. Top: Cardioid co-culture, Bottom: 2D fibronectin re-plating at d8.5 and maintenance or subsequent differentiation.
