## Supplemental Video Legends for "Cardioids reveal self-organizing principles of human cardiogenesis"

**Supplemental Video 1. Effect of ECM Protein Addition on the Formation of 3D Structures During CM Differentiation in Flat Bottom Wells**. Video shows a time-course from d-1 until d6.5 of CM differentiation. Only top half of 96well plate is shown. The plate layout is the following: Columns 1, 2: No ECM addition; Column 3 (10 µg/ml) Vitronectin added in suspension; Column 4: (10 µg/ml) Vitronectin used for coating the wells; Columns 5-8: LN511 added in increasing concentrations: 0.05, 0.1, 0.5 and 2 µg/cm$^2$; Columns 9-12: LN521 added in increasing concentrations: 0.1, 0.5, 1, 5 µg/ml. Video shows one frame per hour.

**Supplemental Video 2. Cardioid at D7.5.** Example of cardiac organoid (WTC line) derived from 5000 hPSCs and treated with 8µM CHIR99021 during induction at d7.5 of differentiation.

**Supplemental Video 3. Calcium-Transients Displayed by a Cardiomyocyte (CM)-cardioid.** Fluo-4 ($Ca^{2+}$ - GFP) loaded CM-cardioid shows the calcium release as it contracts. Scalebar: 200µm.

**Supplemental Video 4. Cardioid Long-Term Culture**. Example of 2-month-old cardioid (WTC line) started from 500 hPSCs and treated with 8µM CHIR99021 during induction.

**Supplemental Video 5. Cavity Formation in Cardioid.** Live imaging of a cardioid derived from a MYH10-GFP reporter line showing the formation of a cavity. Capture starts at d0 (24h). Scalebar: 200 µm.

**Supplemental Video 6. 4.4 kDa Dextran-TAMRA Does Not Get Incorporated Into the Developing Cavities.** 4.4 kDa Dextran-TAMRA was added to the culture medium between hours 64h-90h of cardioid development and did not show signal inside of the cardioid cavities.

**Supplemental Video 7. Single Cardioid Development and Specification.** Time course of a single cardioid developing between hours 16h-178h showing the emergence of cavities during the cardiac mesoderm stage and the successful CM-specification at the end (MYL7-GFP). Scalebar: 1000 µm.

**Supplemental Video 8. Reproducibility of Cardioid development.** Time course of cardioid development between hours 16h-178h showing the reproducible development of cavities and the successful cardiomyocyte (CM)-specification in 48 cardioids (MYL7-GFP expression).
